# Data-driven frame-by-frame analysis of surface swimming and terrestrial walking in the coastal beetle *Phaleria atriceps*

**DOI:** 10.64898/2026.10.22.760847

**Authors:** Kazuki Sakura, Koki Yano, Motoki Nakata, Yasuhiko Chikami

## Abstract

**Background:** Inter-leg coordination of insects changes depending on environmental conditions. Although locomotor changes between terrestrial and aquatic environments have been examined in various insects, relatively little is known about coastal insects living at the boundary between these environments. We therefore compared the inter-leg coordination during terrestrial walking with that during surface swimming in the coastal beetle *Phaleria atriceps*.

**Methods:** Surface swimming was recorded in 17 individuals and terrestrial walking in 10 individuals using high-speed video at 1,000 frames/s. The six leg tips and anterior and posterior body landmarks were tracked frame-by-frame using DeepLabCut, and leg movements were quantified in a body-centered coordinate system. Leg movement periods were estimated from sinusoidal fits to longitudinal leg-tip trajectories. Inter-leg phase relationships were calculated for all 15 leg pairs and integrated to reconstruct a representative six-leg phase organization for each locomotor mode.

**Results:** Leg movement periods were significantly longer during surface swimming than during terrestrial walking for all six legs. During walking, the six-leg phase relationship closely matched an alternating tripod pattern, with a left foreleg (L1), a right middle leg (R2), and a left hind leg (L3) forming one phase group and a right foreleg (R1), a left middle leg (L2), and a right hind leg (R3) forming the opposing group. The same broad grouping was observed in the six-leg phase relationship during surface swimming, although significant differences in relative phase were detected for L1–L2, L1–R2, and R1–R2.

**Conclusions:** *Phaleria atriceps* exhibited six-leg phase relationships broadly consistent with an alternating tripod pattern during both terrestrial walking and surface swimming, whereas leg movement periods and certain inter-leg phase relationships differed between the two locomotor modes. These findings provide a quantitative description of the similarities and differences in inter-leg coordination between terrestrial walking and surface swimming in a coastal beetle. Further mechanical and comparative studies will be needed to clarify the functional and evolutionary significance of these findings.

## Introduction

Insects locomote through diverse environments using coordinated movements of their six legs. Inter-leg coordination varies with locomotor context, speed, and substrate conditions (Weihmann, 2025). One common pattern during terrestrial locomotion is the alternating tripod gait, in which the left foreleg, right middle leg, and left hindleg move approximately in phase, while the right foreleg, left middle leg, and right hindleg form the opposing tripod, and the two tripods move approximately in antiphase (Hughes, 1952). Recent studies have demonstrated that inter-leg coordination differs among different environmental conditions. For example, the ant *Camponotus japonicus* gradually shifts from a tripod pattern to more variable tetrapod-like patterns as the roughness of vertically undulating terrain increases (Zeng et al., 2026). Thus, clarifying how inter-leg coordination differs across different environments is important for understanding insect locomotion.

The air–water interface is mechanically distinct from terrestrial substrates because locomotion at the water surface is affected by hydrodynamic forces (Bush & Hu, 2006). Insects use several locomotion modes at this interface, including walking, rowing and jumping on the water surface (e.g., Bush & Hu, 2006; Hu and Bush 2010; O’Neil et al., 2024) and swimming at the water surface with the body partly immersed, i.e., surface swimming (e.g., Miller, 1972; DuBois & Jander, 1985; Matsuura et al., 2002; Bohn, Thornham & Federle, 2012; Yanoviak & Frederick, 2014). Inter-leg coordination during surface swimming varies among species. In the two-spotted cricket, *Gryllus bimaculatus*, the left and right legs of each pair move approximately in phase during swimming, in contrast to the alternating tripod pattern used during terrestrial walking (Matsuura et al., 2002; Naniwa et al., 2020). In contrast, the ant *Camponotus schmitzi* exhibits an alternating tripod gait during surface swimming and terrestrial walking, although the stepping frequency and inter-leg phase relationships are different between the two locomotor modes (Bohn, Thornham & Federle, 2012). Similarly, the canopy ants *Pachycondyla spp*. and *Odontomachus bauri* use a modified alternating tripod gait during surface swimming (Yanoviak & Frederick, 2014).

Coastal insects provide an informative system examining locomotion at the boundary between terrestrial and aquatic environments. The darkling beetle *Phaleria atriceps* (Lewis, 1894) inhabits near the tideline on sandy beaches in Japan and has been collected beneath seaweed (Lewis 1894; Grimm & Lillig, 2020; Chiba, Ichinose & Karasawa, 2026). However, quantitative information on inter-leg coordination during surface swimming remains limited for coastal terrestrial beetles, including *P. atriceps*. Here, we compared leg movement periods and inter-leg phase relationships between surface swimming and terrestrial walking in *P. atriceps* using frame-by-frame tracking of the six leg tips.

## Materials & Methods

### Insects

We collected 17 adult *Phaleria atriceps* at Nanane Coast in Toyohashi City, Aichi Prefecture, Japan (34°39′48″N, 137°25′03″E), on 29 June 2022, and 10 adults at the same location on 28 August 2025 (Figure 1A, B). The 17 individuals collected in 2022 were used for surface swimming recordings, whereas the 10 individuals collected in 2025 were used for terrestrial walking recordings. The beetles were maintained under a 16 h light:8 h dark photoperiod at 25.0 ± 1.0 °C and were fed insect jelly (Roine, Japan). Behavioral recordings were conducted 1–5 days after collection.

**Figure 1.**
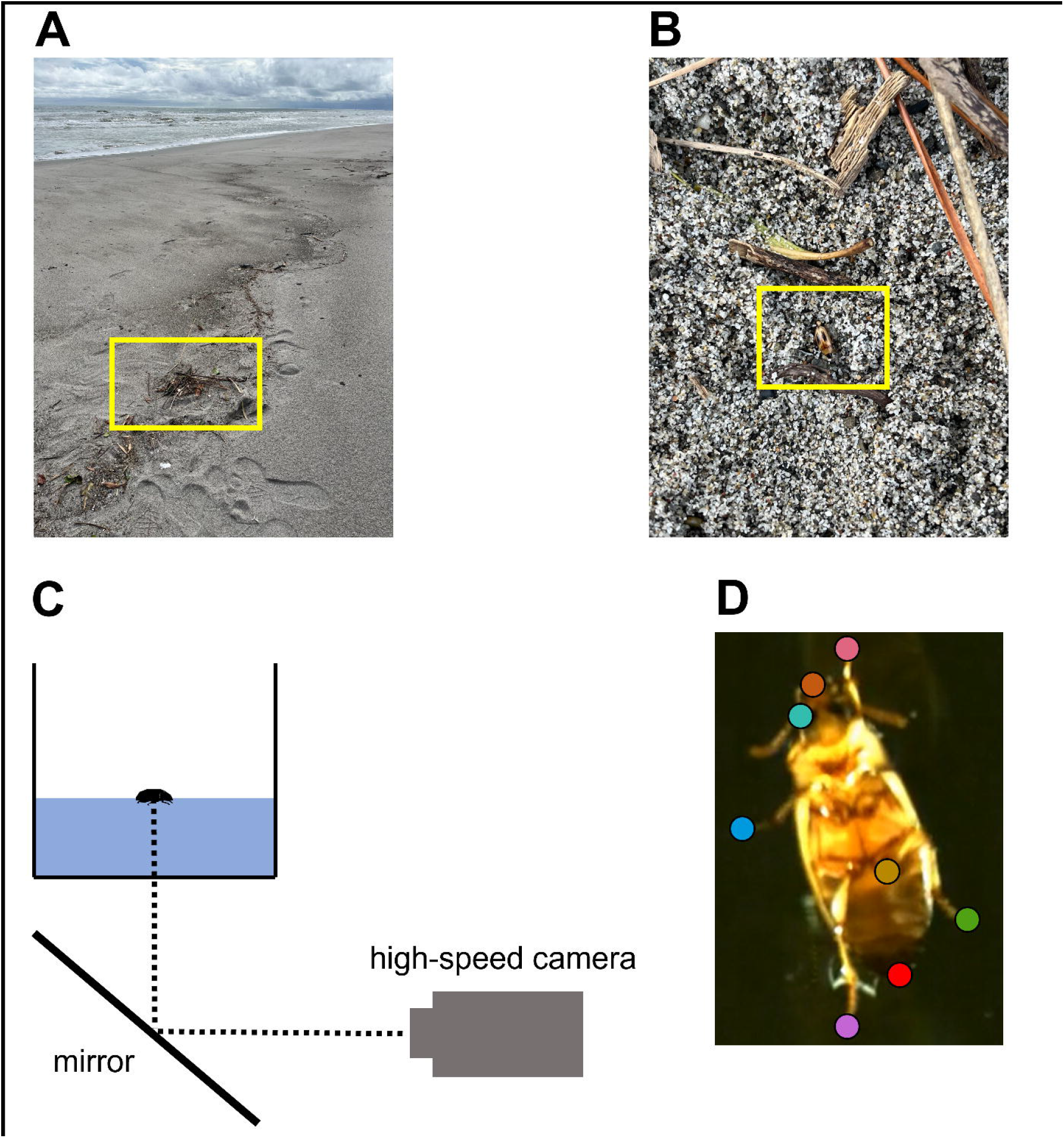
Collection site, experimental setup, and DeepLabCut landmark detection during surface swimming in *Phaleria atriceps*. (A) Collection site on the coast, showing stranded seaweed under which *P. atriceps* were found. The collection area is indicated by the yellow box. (B) *P. atriceps* exposed after the stranded seaweed was moved. The beetle is indicated by the yellow box. (C) Illustration of the experimental setup used for recording of surface swimming. (D) Representative frame from a swimming video, showing the positions of the six leg tips and the anterior and posterior body landmarks detected using DeepLabCut (colored circles). The photographs in (A), (B) and the video frame in (D) were captured by Kazuki Sakura.

### Recording surface swimming and terrestrial walking

For surface swimming recording, each of the 17 *P. atriceps* individuals was placed separately in a tank (30 × 30 × 30 cm) containing water to a depth of 10 cm. We defined swimming as directional locomotion through the water. During swimming, the body was positioned near the air–water interface while the legs were submerged in the water. Therefore, we refer to this behavior as surface swimming. A mirror was placed beneath the tank, and the image reflected in the mirror was recorded at 1,000 frames/s using a high-speed camera (FASTCAM SA-Z, Photron, Japan) (Figure 1C). The surface swimming dataset consisted of 17 videos, each obtained from a different individual.

For recording terrestrial walking, each of the 10 *P. atriceps* individuals was placed separately on a paper and recorded from above at 1,000 frames/s using the same high-speed camera. The terrestrial walking dataset consisted of 10 videos, each obtained from a different individual.

### Frame-by-frame tracking of leg movements

Swimming videos were analyzed using DeepLabCut version 3.0.1 with the PyTorch engine (Mathis et al., 2018; Nath et al., 2019). A single-animal pose-estimation project was constructed using all 17 swimming videos. Eight landmarks were tracked: the tips of the left foreleg (L1), left middle leg (L2), left hindleg (L3), right foreleg (R1), right middle leg (R2), and right hindleg (R3), together with the anterior and posterior ends of the body (Figure 1D). Fifty frames from each video were manually annotated, resulting in 850 labeled frames in total. The pose-estimation network used a ResNet-50 backbone, with 95% of the labeled frames assigned to the training set. The network was trained for 200 epochs using the AdamW optimizer with a batch size of 8. After training, the network was applied to all swimming videos to obtain frame-by-frame coordinates.

Walking videos were analyzed using a separate single-animal DeepLabCut project. The same eight anatomical landmarks were tracked. Fifty frames were randomly selected from each of the 10 walking videos and manually annotated, yielding 500 labeled frames in total. A ResNet-50 backbone was used, and the network was trained for 200 epochs. The trained model was then applied to all 10 walking videos to obtain frame-by-frame coordinates. The other training settings were same as those used for analysis for swimming videos.

### Body-centered leg trajectories

All subsequent analyses were performed in R version 4.6.1. For each frame, the midpoint between the anterior and posterior body landmarks was defined as the body center, and the line from the posterior to the anterior landmark defined the longitudinal body axis. Coordinates were translated so that the body center was located at the origin and rotated so that the longitudinal body axis was aligned with the longitudinal coordinate axis. Coordinates were normalized by the median anterior–posterior body length of each recording. Subsequent analyses were performed using frames in which the DeepLabCut likelihood values of the leg tip and both body landmarks were at least 0.2.

### Estimation of leg movement period

Periodic leg movement was quantified separately for each leg in each recording. The longitudinal trajectory *η*_*l*_ of leg *l* was fitted with

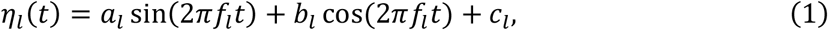

where *f*_*l*_ denotes the frequency determined for leg *l* and *t* denotes the time, *a*_*l*_ and *b*_*l*_ are the sine and cosine coefficients, respectively, and *c*_*l*_ is the constant offset. Candidate frequencies ranging from 0.5 to 20 Hz were evaluated. At each candidate frequency, *a*_*l*_, *b*_*l*_ and *c*_*l*_ were fitted to the longitudinal trajectory, and goodness of fit was quantified using *R*^2^. Local maxima of *R*^2^ across the tested frequency range were identified as candidate movement frequencies. Only trajectories for which the best-fitting sinusoid had *R*^2^ ≥ 0.2 were retained for subsequent analyses.

Because the movement frequency of a given leg was expected to be broadly similar among individuals performing the same locomotor mode, frequency estimates from the other recordings were used to distinguish the fundamental movement frequency from harmonic components. For each leg in each recording, a reference frequency was calculated as the median of the frequencies yielding the highest *R*^2^ for the same leg in all other recordings of the same locomotor mode. The candidate movement frequency with the highest *R*^2^ was retained when it was consistent with this reference frequency.

In some trajectories, the highest *R*^2^ occurred at approximately twice the reference frequency, suggesting that the second harmonic, rather than the fundamental movement cycle, produced the strongest sinusoidal fit. In such cases, the lower-frequency component was accepted as the fundamental frequency only when a distinct local maximum in *R*^2^ is present near one-half of the frequency associated with the highest *R*^2^. Detailed criteria for identifying candidate movement frequencies and second harmonics are provided in Supplementary Methods.

### Estimation of pairwise phase relationships between legs

For each leg, the longitudinal trajectory was refitted using Eq. (1) with the frequency fixed at the value estimated in the previous analysis. The phase offset *ϕ*_*l*_ for leg *l* was calculated from the fitted sine and cosine coefficients as

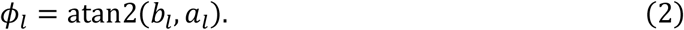

The phase *θ*_*l*_ of leg *l* at time *t* was then expressed as

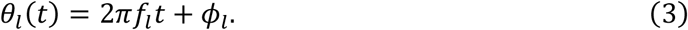

Because the frequency was estimated independently for each leg, the two legs within a pair were not assumed to move at the same frequency. Consequently, their relative phase could vary over time. We therefore evaluated the relative phase at the midpoint of the interval bounded by the first and last frames in which both legs simultaneously met the tracking quality criterion. For each leg, this criterion required the DeepLabCut likelihood values for the leg tip and both body landmarks to be at least 0.2. For a pair consisting of leg 1 and leg 2, the representative relative phase Δ*ϕ* was calculated as

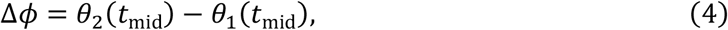

where *t*_mid_ is the midpoint of this interval.

### Estimation of the six-leg phase relationship

For each locomotor mode, we estimated representative phase relationships among the six legs from the pairwise phase relationships obtained for the 15 leg pairs in the previous subsection. Let 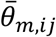 denote the circular mean phase of leg *j* with respect to leg *i* in locomotor mode *m*, and let *β*_m,l_ denotes the phase of leg *l* among the six legs. The six-leg phases were estimated by minimizing

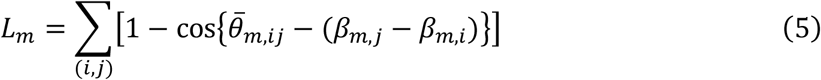

across all 15 leg pairs. Because phases between two legs were relative values, the phase of L1 was fixed at 0° as a reference for the other legs.

### Statistical analysis

The linear mixed-effects model was fitted using the “nlme” package (version 3.1-169; Pinheiro, Bates & R Core Team, 2026). Estimated marginal means and within-leg comparisons between swimming and walking were obtained using the “emmeans” package (version 2.0.4; Lenth & Piaskowski, 2026).

#### Comparison of leg movement period

Leg movement periods were log-transformed before statistical analysis. Because up to six leg measurements were obtained from the same individual, we used a linear mixed-effects model with locomotor mode, anatomical leg, and their interaction as fixed effects and the individual as a random intercept:

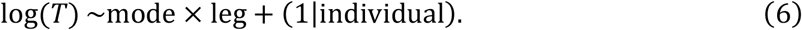

Because residual variability appeared to differ between swimming and walking, the model allowed separate residual variances for the two locomotor modes. Swimming and walking were compared separately for each leg, with *P*-values adjusted across the six comparisons using the Holm procedure.

#### Comparison of pairwise phase relationships between swimming and walking

Swimming and walking were compared separately for each pairwise phase relationship among the 15 leg pairs using an exact permutation test. For each pair, the difference between the circular mean phases of swimming and walking was quantified using

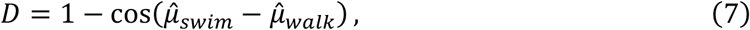

where 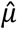 denotes the circular mean. The statistic *D* equals 0 when the two mean directions are identical and increases as their angular separation increases. Statistical significance was determined by enumerating all possible reallocations of the observed phase values between swimming and walking while preserving the observed sample sizes for that leg pair. The resulting *P*-values for the 15 leg pairs were adjusted using the Holm procedure.

#### Comparison of six-leg phase relationships between swimming and walking

Because the phase of L1 was set to 0° separately when estimating the swimming and walking six-leg phase relationships, they were aligned before calculating the phase differences of the legs. For each leg *l*, the angular difference between swimming and walking was first calculated as

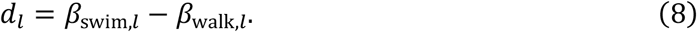

The common rotation between the two six-leg phase relationships was defined as the circular mean of the six angular differences,

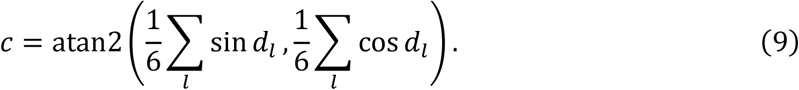

The six-leg phase relationship during swimming was then rotated by −*c*, and the phase difference of each leg between walking and swimming was calculated as

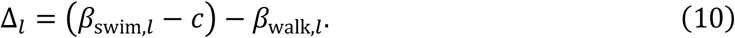

Positive values of Δ_*l*_ indicate that the phase of a given leg was relatively advanced during swimming, whereas negative values indicate a relative phase delay.

Confidence intervals for the estimated phase differences were evaluated by bootstrap resampling of individuals. For each bootstrap replicate, the circular mean phase for each leg pair, the six-leg phase relationship, the alignment between locomotor modes, and Δ_*l*_ were recalculated.

A total of 5,000 bootstrap replicates were obtained. The 95% confidence interval for each leg was calculated from the bootstrap distribution.

## Results

### Sinusoidal fitting of leg trajectories

To quantify the periodicity of leg movements and identify the fundamental movement frequencies used in the subsequent analyses of leg movement period and inter-leg phase relationships, we fitted a sinusoidal model to the longitudinal trajectories of the leg tips. 60 of 102 swimming trajectories and all 60 walking trajectories satisfied the fit quality threshold *R*^2^ ≥ 0.2.

Among the 60 swimming trajectories, the frequency associated with the highest *R*^2^ value was consistent with the reference frequency derived from the same leg in other swimming recordings in 45 trajectories and was directly selected as the fundamental frequency (Figure 2A, E, I). In four trajectories, the frequency associated with the highest *R*^2^ value occurred near twice the reference frequency, and a distinct local peak near half of the dominant frequency was selected as the fundamental frequency (Figure 2B, F, J). The remaining 11 trajectories were excluded because their frequency structure was inconsistent with the reference frequency (Figure 2C, G, K). In walking, the frequency associated with the highest *R*^2^ value was consistent with the reference frequency in all 60 trajectories and was directly selected as the fundamental frequency (Figure 2D, H, L).

**Figure 2.**
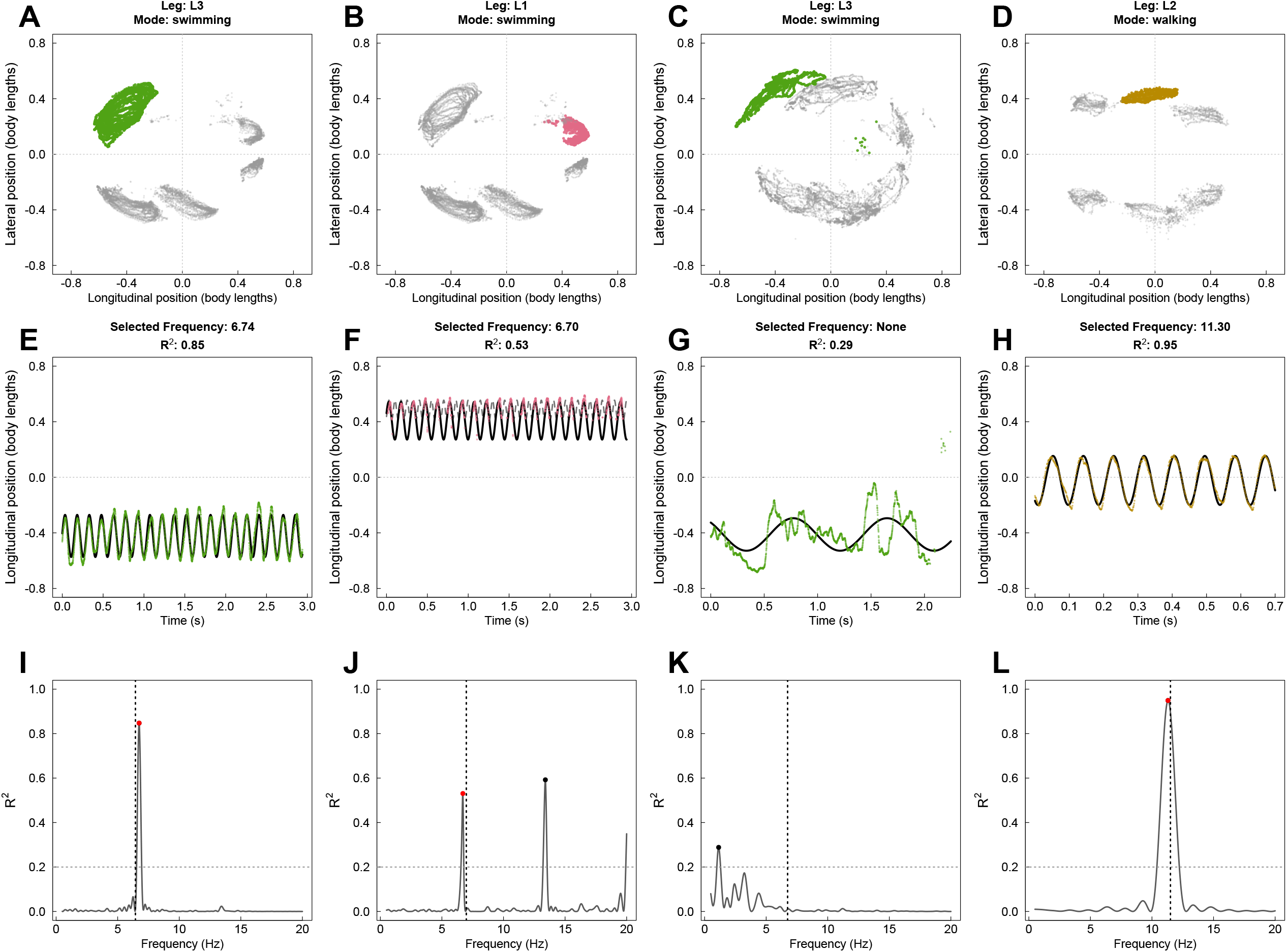
Representative sinusoidal fitting and selection of fundamental leg movement frequency in *Phaleria atriceps*. (A–D) Body-centered trajectories of the six leg tips for representative recordings. The leg used for frequency estimation is shown in color, and the other legs are shown in gray. Longitudinal and lateral positions are expressed relative to body length. Only frames in which the DeepLabCut likelihood values for the corresponding leg tip and both body landmarks were at least 0.2 are shown. (E–H) Time series of longitudinal position of the leg used for frequency estimation. Colored points show the observed trajectory, and the black solid line shows the sinusoidal fit at the selected frequency in (E), (F), and (H). In (G), no fundamental frequency was selected, and the black solid curve instead shows the fit at the frequency with the highest *R*^2^ value. (I–L) *R*^2^ obtained by fitting sinusoidal models. The vertical dotted line indicates the leave-one-out reference frequency derived from the same leg in the other recordings, and the horizontal dotted line indicates the fit quality threshold *R*^2^ ≥ 0.2. Black points indicate the dominant *R*^2^ peak, and red points indicate the frequency selected as the fundamental frequency. (A, E, I) Example of a swimming trajectory in which the dominant peak was consistent with the reference frequency and was directly selected. (B, F, J) Example of a swimming trajectory in which the dominant peak occurred near twice the reference frequency and a lower-frequency local peak near the reference frequency was selected as the fundamental frequency; the gray dashed curve in F shows the sinusoidal fit at the frequency of the dominant peak. (C, G, K) Example of a swimming trajectory excluded because the dominant frequency structure was inconsistent with the reference frequency; no fundamental frequency was selected. (D, H, L) Example of a walking trajectory in which the dominant peak was consistent with the reference frequency and was directly selected.

For trajectories in which the frequency associated with the highest *R*^2^ value was directly selected, the median *R*^2^ of the sinusoidal fit was 0.515 (from 0.210 to 0.847) during swimming and 0.867 (from 0.312 to 0.950) during walking. The final selected fundamental frequencies were centered around 7 Hz during swimming (mean = 7.00 Hz; median = 6.94 Hz) and 11 Hz during walking (mean = 11.46 Hz; median = 11.38 Hz).

### Leg movement period is shorter during terrestrial walking than surface swimming

We next examined whether leg movement period differed between surface swimming and terrestrial walking. For each retained leg trajectory, movement period was calculated from the selected fundamental frequency as 1⁄*f*. The linear mixed effects model revealed a significant effect of locomotor mode, i.e., swimming vs walking, on log-transformed period (*F*_1,20_ = 298.86, *P* = 1.71 × 10^-13^), whereas neither the effect of leg (*F*_S,77_ = 0.75, *P* = 0.590) nor the interaction between locomotor mode and leg (*F*_S.77_ = 1.47, *P* = 0.211) was significant. Comparisons between swimming and walking for each leg showed that leg movement periods were significantly shorter during walking than during swimming for all six legs (Holm adjusted *P* < 0.001 for all comparisons; Figure 3). Across the six legs, mean leg movement periods ranged from 0.140 to 0.147 s during swimming and from 0.0874 to 0.0879 s during walking.

**Figure 3.**
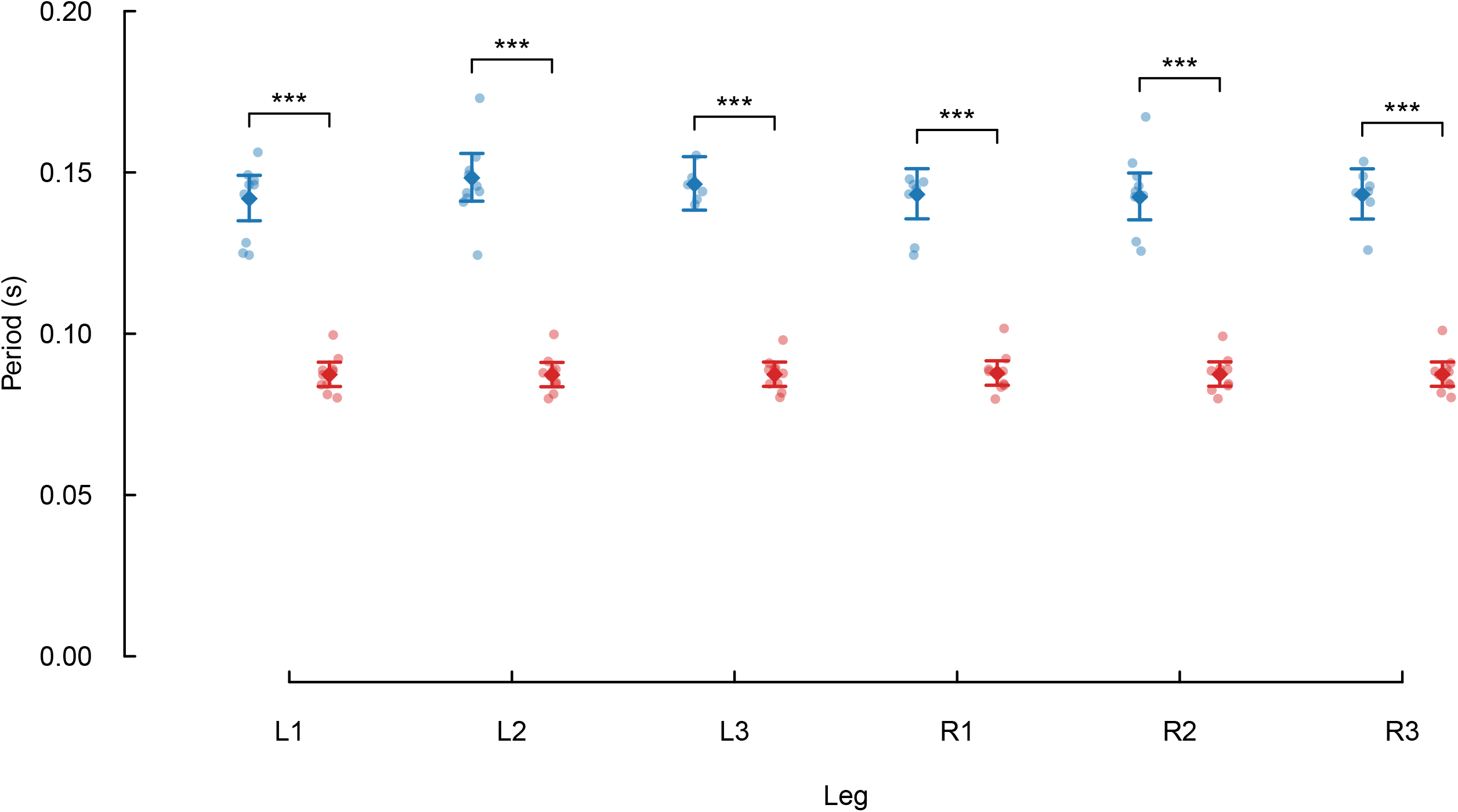
Leg movement period during surface swimming and terrestrial walking in *Phaleria atriceps*. Leg movement period was calculated from the fundamental frequency for each leg trajectory selected by the sinusoidal fitting. Small points show periods for each of the six legs during surface swimming (blue) and terrestrial walking (red). Diamonds indicate model-estimated periods for each locomotor mode and leg. The periods are transformed from the log scale. Error bars indicate 95% confidence intervals. Periods were analyzed using a linear mixed-effects model with locomotor mode, leg, and their interaction as fixed effects, individual as a random intercept, and separate residual variances for swimming and walking. Brackets indicate comparisons between swimming and walking within each leg. *P*-values for the six within-leg comparisons were adjusted using the Holm method; \*\*\**P* < 0.001.

### Phase relationships among legs differ between surface swimming and terrestrial walking

We next examined whether inter-leg phase relationships differ between surface swimming and terrestrial walking. Pairwise phase relationships were quantified for each of the 15 leg pairs and compared between the two locomotor modes. Following Holm correction for multiple comparisons, significant differences between swimming and walking were detected for three pairs: L1–L2, L1–R2, and R1–R2 (Fig. S1). The circular mean phase for L1–L2 was 122.1° during swimming and 189.3° during walking, corresponding to an absolute angular difference of 67.2° (*P* = 0.0154, Holm correction). For L1–R2, the mean phase was 302.2° during swimming and 10.6° during walking, with an absolute angular difference of 68.3° (*P* = 0.000771, Holm correction). For R1–R2, the mean phase was 129.4° during swimming and 187.5° during walking, with an absolute angular difference of 58.1° (*P* = 0.00466, Holm correction). No significant differences between locomotor modes were detected for the remaining 12 pairs (Fig. S1).

We then estimated the six-leg phase relationships from the pairwise phase relationships of the 15 leg pairs separately for each locomotor mode. The phase of L1 was fixed at 0° as a reference for both walking and swimming. During walking, R2 and L3 were positioned at 10.6° and 6.8°, respectively, whereas R1, L2, and R3 were positioned at 183.0°, 189.2°, and 187.1°, respectively (Figure 4A). Thus, the six-leg phase relationship during walking closely corresponded to an alternating tripod pattern, with L1, R2, and L3 forming one phase group and R1, L2, and R3 forming the opposing group approximately 180° away (Figure 4A).

**Figure 4.**
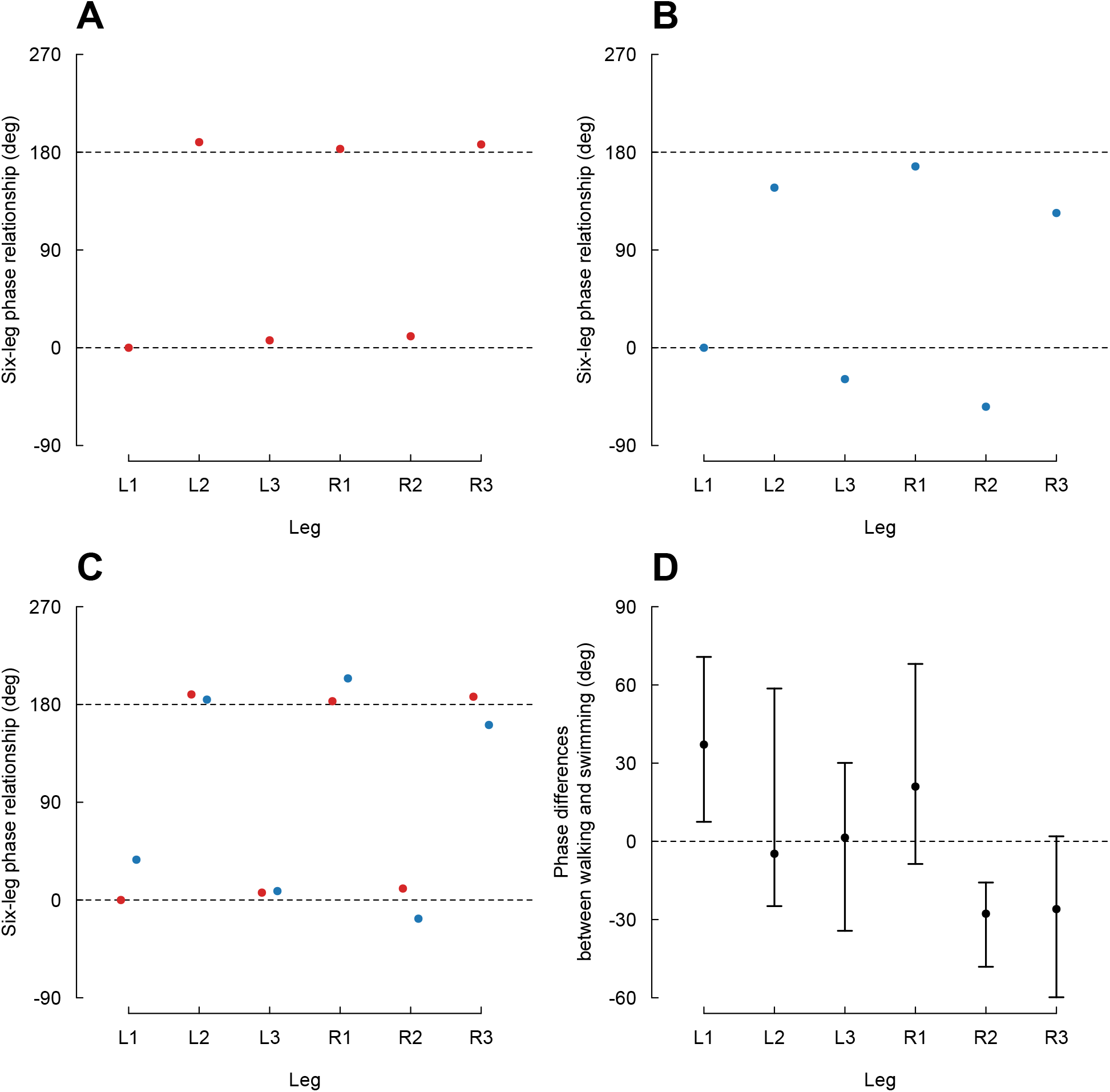
Six-leg phase relationships during terrestrial walking and surface swimming in *Phaleria atriceps*. (A) Six-leg phase relationships during terrestrial walking (red), with the phase of L1 fixed at 0°. (B) Six-leg phase relationships during surface swimming (blue), independently estimated with the phase of L1 fixed at 0°. The horizontal dotted lines in (A) and (B) indicate 0° and 180°. (C) Comparison of the six-leg phase relationships during walking (red) and swimming (blue) after alignment. The horizontal dotted lines show 0° and 180°. (D) Phase differences between terrestrial walking and surface swimming. Points indicate the estimated phase differences, and error bars indicate 95% bootstrap confidence intervals based on 5,000 bootstrap replicates. The horizontal dotted lines show 0°.

During surface swimming, R2 and L3 were positioned at −54.3° and −28.9°, respectively, whereas R1, L2, and R3 were positioned at 166.9°, 147.4°, and 124.0°, respectively, relative to L1 (0°; Figure 4B). The six-leg phase relationship during swimming was also broadly consistent with an alternating tripod pattern, although the phases of individual legs deviated from the ideal values of 0° and 180°.

To compare the six-leg phase relationships between the two locomotor modes, we then aligned the six-leg phase relationship of swimming with that of walking by applying a common angular rotation to phases of all six legs during swimming (Figure 4C). After alignment, the six-leg phase relationship during swimming, L1, R2, and L3 were positioned at 37.1°, −17.2°, and 8.2° respectively, whereas R1, L2, and R3 were positioned at 204.0°, 184.5°, and 161.1°, respectively (Figure 4C).

We finally quantified the relative phase difference of each leg between walking and swimming (Figure 4D). The estimated differences were +37.1° for L1, −4.8° for L2, +1.4° for L3, +21.0° for R1, −27.8° for R2, and −26.0° for R3 (Figure 4D). The 95% bootstrap confidence interval did not include zero for L1 (7.5° to 70.7°) or R2 (−48.1° to −15.8°), whereas the intervals for the other four legs included zero. Overall, six-leg phase relationships during walking and swimming are broadly consistent with an alternating tripod pattern, but there are relative phase differences between swimming and walking: L1 showed relative phase advance, R2 showed relative phase delay.

## Discussion

The present study quantitatively compared leg movement periods and inter-leg phase relationships between terrestrial walking and surface swimming in the coastal darkling beetle *Phaleria atriceps*. The six-leg phase relationships during both locomotor modes were broadly consistent with an alternating tripod pattern, with a left foreleg (L1), a right middle leg (R2), and a left hind leg (L3) forming one phase group and a right foreleg (R1), a left middle leg (L2), and a right hind leg (R3) forming the opposing group. However, leg movement periods were longer during surface swimming, and L1 showed relative phase advance and R2 showed relative phase delay during surface swimming compared with terrestrial walking.

A similar result has been reported in the pitcher plant ant *Camponotus schmitzi*. During both terrestrial running and surface swimming, the same two sets of legs form alternating tripod patterns, but the phase delays among legs within each tripod are larger during swimming (Bohn, Thornham & Federle, 2012). In contrast, other insects show inter-leg phase relationships during surface swimming different from those during terrestrial walking. In the two-spotted cricket, *Gryllus bimaculatus*, for example, while its legs move in an alternating tripod pattern during terrestrial walking, the left and right leg pairs move approximately in phase in a breaststroke-like pattern during surface swimming (Matsuura et al., 2002; Naniwa et al., 2020). The mantis *Sphodromantis lineola* swims using alternating strokes of the middle and hind leg pairs, with the left and right legs of each pair moving synchronously (Miller, 1972). These examples demonstrate the diversity of inter-leg phase relationships during surface swimming. This diversity may reflect the following two factors. First, different inter-leg coordination may be mechanically advantageous under different combinations of body morphology and environmental conditions. For example, experiments with an amphibious turtle-inspired robot showed that locomotor efficiency depended on the combination of gait, limb shape, and environmental medium (Baines et al., 2022). Similarly, studies of aquatic propulsion have shown that swimming performance can depend on whether legs move synchronously or alternately, as well as on the phase lag among appendages (Whittlesey, 2011; Ford & Santhanakrishnan, 2021). Thus, there may be no single inter-leg coordination that is optimal for aquatic locomotion across taxa. Second, the inter-leg coordination expressed in water may also be influenced by phylogenetic history. In a broad comparative analysis of spiders, Stratton et al. (2004) found that phylogeny was often a stronger predictor of rowing behavior than current association with aquatic habitats. In *P. atriceps*, one possibility is that the inter-leg phase relationship during surface swimming may be mechanically advantageous for locomotion at the water surface. In particular, the phase relationships between forelegs and middle legs could influence the temporal distribution of hydrodynamic forces and affect propulsion and stability. Alternatively, the inter-leg phase relationship during swimming of *P. atriceps* may represent an ancestral pattern maintained from its ancestor species, regardless of whether it is mechanically optimal for surface swimming. Further studies combining measurements of swimming mechanics with comparisons among related beetle species will be needed to evaluate these possibilities.

Leg movement period also significantly differed between the two locomotor modes. A similar result was reported in *C. schmitzi*, in which leg movement frequency was lower during swimming than during terrestrial running (Bohn et al., 2012). Bohn et al. (2012) suggested that differences between swimming and running kinematics may partly arise from the different physical constraints imposed by the two environments. In particular, leg protraction through water may be slower than through air because of hydrodynamic resistance, while terrestrial locomotion imposes constraints associated with ground contact and body support. Similar mechanical differences may contribute to the longer leg movement periods observed during swimming in *P. atriceps*. For example, hydrodynamic resistance may slow forward leg movements, thereby prolonging leg movement cycle. Direct measurements of leg forces and the surrounding flow will be needed to determine how these physical factors contribute to the observed difference in leg movement period.

Previous studies have characterized inter-leg phase relationships during aquatic locomotion primarily using event-based approaches. Several studies estimated inter-leg coordination from the timing of power and recovery strokes (Bowdan, 1978; Matsuura et al., 2002; Bohn et al., 2012; Naniwa et al., 2020; Ruszczyk et al. 2021). In the present study, however, the trajectory data contained noise and missing values, particularly around transitions from power to recovery strokes in the forelegs during surface swimming (Figure 2F). Because these transitions could not be reliably identified, we could not apply an event-based approach. We therefore estimated phase from leg-tip trajectories using sinusoidal fitting. We then quantified phase relationships for all leg pairs and integrated them into a representative six-leg phase relationship. This approach allowed us to assess not only whether walking and swimming shared the broad alternating tripod pattern, but also how inter-leg coordination differed between terrestrial walking and surface swimming.

## Conclusions

Insect inter-leg coordination differs depending on environmental conditions. Although differences in locomotion modes between terrestrial and aquatic environments have been examined in various insects, much less is known about coastal insects living at the boundary between these environments. Here, we focused on the coastal beetle *Phaleria atriceps* (Lewis, 1894) and compared its leg movement periods and inter-leg phase relationships during surface swimming with those during terrestrial walking. The broad alternating tripod pattern was observed during both locomotion modes, but leg movement periods were longer during surface swimming and several relative phase relationships differed. These findings provide a quantitative description of how locomotor coordination differs in a coastal insect. In future, comparative studies and more detailed mechanical studies will contribute to understanding how this difference has evolved.

## Acknowledgements

We would like to thank Dr. Teruyuki Niimi for providing the experimental facilities and for fruitful discussions. We also thank Dr. Hiroyo Nishide for her assistance with the DeepLabCut computations performed on the supercomputer at the National Institute for Basic Biology. We are grateful to Mr. Toshiyuki Sazi for preparing the experimental setup. We used ChatGPT (GPT-6; OpenAI) for proofreading the manuscript and R scripts used for data analysis, with the aim of improving clarity and identifying errors.

## Figure legends

Figure S1 Relative phase relationships during surface swimming and terrestrial walking. Circular plots show the relative phase for each of the 15 possible pairs of legs during surface swimming (blue) and terrestrial walking (red). In each pair name, the first leg was used as the reference; for example, for L1–L2, relative phase was defined as the phase of L2 minus that of L1. Each point represents one individual. Swimming and walking values are plotted at different radial positions only to improve visibility. Arrows indicate the circular mean phase for each locomotor mode, with arrow length proportional to the resultant length, which describes the concentration of phase values among individuals. Differences in circular mean phase between swimming and walking were tested separately for each leg pair using an exact permutation test, and *P*-values for the 15 comparisons were adjusted using the Holm method. \**P* < 0.05, \*\**P* < 0.01, \*\*\**P* < 0.001.

## References

Baines R, Patiballa SK, Booth J, Ramirez L, Sipple T, Garcia A, Fish F, Kramer-Bottiglio R.2022. Multi-environment robotic transitions through adaptive morphogenesis. Nature 610:283–289 DOI 10.1038/s41586-022-05188-w.

Bohn HF, Thornham DG, Federle W. 2012. Ants swimming in pitcher plants: kinematics of aquatic and terrestrial locomotion in Camponotus schmitzi. Journal of Comparative Physiology A 198:465–476 DOI 10.1007/s00359-012-0723-4.

Bowdan E. 1978. Walking and rowing in the water strider, Gerris remigis. I. A cinematographic analysis of walking. Journal of Comparative Physiology A 123:43–49 DOI 10.1007/BF00657342.

Bush JWM, Hu DL. 2006. Walking on water: biolocomotion at the interface. Annual Review of Fluid Mechanics 38:339–369.

Chiba H, Ichinose A, Karasawa S. 2026. Zonation of soil-dwelling invertebrates on a natural sandy beach, Tottori, western Japan. Entomological Science 29:e70019 DOI 10.1111/ens.70019.

DuBois MB, Jander R. 1985. Leg coordination and swimming in an ant, Camponotus americanus. Physiological Entomology 10:267–270 DOI 10.1111/j.1365-3032.1985.tb00046.x.

Ford MP, Santhanakrishnan A. 2021. On the role of phase lag in multi-appendage metachronal swimming of euphausiids. Bioinspiration & Biomimetics 16:066007 DOI 10.1088/1748-3190/abc930.

Franklin R, Jander R, Ele K. 1977. The coordination, mechanics and evolution of swimming by a grasshopper, Melanoplus differentialis (Orthoptera: Acrididae). Journal of the Kansas Entomological Society 50:189–199.

Grimm R, Lillig M. 2020. Description of a new Phaleria Latreille, 1802 from Bali and a new synonym of Phaleria atriceps (Lewis, 1894) (Insecta: Coleoptera: Tenebrionidae: Diaperinae: Phaleriini). Vernate 39:301–305.

Hu DL, Bush JWM. 2010. The hydrodynamics of water-walking arthropods. Journal of Fluid Mechanics 644:5–33 DOI 10.1017/S0022112009992205.

Hughes GM. 1952. The co-ordination of insect movements. I. The walking movements of insects.Journal of Experimental Biology 29:267–285 DOI 10.1242/jeb.29.2.267.

Lenth R, Piaskowski J (2026). emmeans: Estimated marginal means, aka least-squares means. R package version 2.0.4. 10.32614/CRAN.package.emmeans

Lewis G. 1894. On the Tenebrionidae of Japan. Annals and Magazine of Natural History, Series 6 13:377–400.

Mathis A, Mamidanna P, Cury KM, Abe T, Murthy VN, Mathis MW, Bethge M. 2018.DeepLabCut: markerless pose estimation of user-defined body parts with deep learning.Nature Neuroscience 21:1281–1289 DOI 10.1038/s41593-018-0209-y.

Matsuura T, Kanou M, Yamaguchi T. 2002. Motor program initiation and selection in crickets, with special reference to swimming and flying behavior. Journal of Comparative Physiology A 187:987–995 DOI 10.1007/s00359-001-0269-3.

Miller PL. 1972. Swimming in mantids. Journal of Entomology, Series A: General Entomology46:91–97 DOI 10.1111/j.1365-3032.1972.tb00113.x.

Naniwa K, Sugimoto Y, Osuka K, Aonuma H. 2020. Novel method for analyzing flexible locomotion patterns of animals by using polar histogram. Journal of Robotics and Mechatronics 32:812–821 DOI 10.20965/jrm.2020.p0812.

Nath T, Mathis A, Chen AC, Patel A, Bethge M, Mathis MW. 2019. Using DeepLabCut for 3D markerless pose estimation across species and behaviors. Nature Protocols 14:2152–2176 DOI 10.1038/s41596-019-0176-0.

O’Neil JN, Yung KL, Difini G, Walker H, Bhamla MS. 2024. Tiny amphibious insects use tripod gait for traversal on land, water, and duckweed. Integrative and Comparative Biology 64:1044–1054 DOI 10.1093/icb/icae078.

Pinheiro J, Bates D R Core Team (2026). nlme: Linear and nonlinear mixed effects models. R package version 3. 1–169. 10.32614/CRAN.package.nlme

R Core Team (2026). R: A language and environment for statistical computing. R Foundation for Statistical Computing, Vienna, Austria. https://www.R-project.org/

Ruszczyk M, Webster DR, Yen J. 2021. Dual phase-shifted ipsilateral metachrony in Americamysis bahia. Integrative and Comparative Biology 61:1644–1657 DOI 10.1093/icb/icab119.

Stratton GE, Suter RB, Miller PR. 2004. Evolution of water surface locomotion by spiders: a comparative approach. Biological Journal of the Linnean Society 81:63–78 DOI 10.1111/j.1095-8312.2004.00269.x.

Weihmann T. 2025. Interactions of morphology and leg-driven locomotor behaviour in insects. In: Betz O, ed. Insect Ecomorphology: Linking Functional Insect Morphology to Ecology and Evolution. Academic Press, 191–225 DOI 10.1016/B978-0-443-18544-1.00017-X.

Whittlesey RW. 2011. Wake-based unsteady modeling of the aquatic beetle Dytiscus marginalis.Journal of Theoretical Biology 291:14–21 DOI 10.1016/j.jtbi.2011.08.005.

Yanoviak SP, Frederick DN. 2014. Water surface locomotion in tropical canopy ants. Journal of Experimental Biology 217:2163–2170 DOI 10.1242/jeb.101600.

Zeng M, Mao L, Meng C, Ding Y, Fu H, Ansari M, Luo Z, Yu J, An S. 2026. Camponotus japonicus locomotion on vertically undulating mesoscale rough terrain. Journal of Experimental Biology 229:jeb251655 DOI 10.1242/jeb.251655.

