## Supplementary figures and images for "Data-driven frame-by-frame analysis of surface swimming and terrestrial walking in the coastal beetle *Phaleria atriceps*"

### Supplemental Figure 1

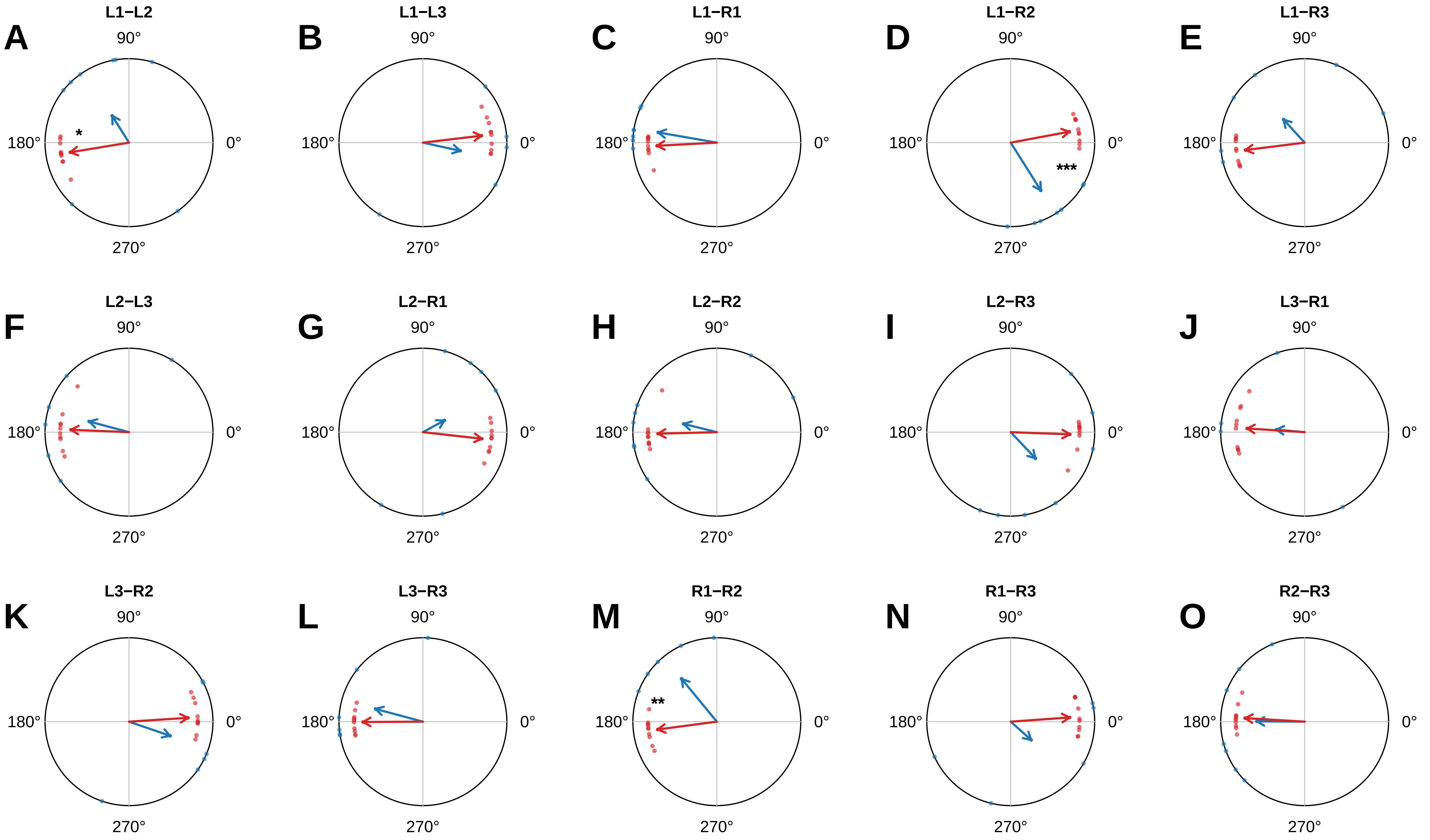
