## Supplemental Methods for "Data-driven frame-by-frame analysis of surface swimming and terrestrial walking in the coastal beetle *Phaleria atriceps*": 01_coordinate_transform.R

```
RAW_DIR = "raw_data"
OUTPUT_DIR = file.path("scratch_results", "coordinate_transform")
RECORDING_DIR = file.path(OUTPUT_DIR, "recordings")
PLOT_DIR = file.path(OUTPUT_DIR, "plots")
FPS = 1000
LEGS = c("L1", "L2", "L3", "R1", "R2", "R3")
BODYPARTS = c(LEGS, "head", "tail")
COORD = c("x", "y", "likelihood")
LEG_COLOR = setNames(hcl.colors(length(LEGS), "Dark 3"), LEGS)
LIKELIHOOD_THRESHOLD = 0.2

dir.create(OUTPUT_DIR, recursive = TRUE, showWarnings = FALSE)
dir.create(RECORDING_DIR, recursive = TRUE, showWarnings = FALSE)
dir.create(PLOT_DIR, recursive = TRUE, showWarnings = FALSE)

make_file_info = function(path) {
    file_info = data.frame()
    file_name = tools::file_path_sans_ext(basename(path))
    all_file_info = regmatches(file_name, regexec("^(swim|walk)_([0-9]+)", file_name))[[1]]
    mode = all_file_info[2]
    number = as.integer(all_file_info[3])
    file_info = data.frame(id = paste0(mode, "_", number), mode = mode, stringsAsFactors = FALSE)
    return(file_info)
}

read_dlc = function(path) {
    raw = read.csv(path, header = FALSE, stringsAsFactors = FALSE,
                   check.names = FALSE, na.strings = c("", "NA", "NaN", "nan"))
    data = vector("list", length(BODYPARTS)*length(COORD))
    i = 1
    for (part in BODYPARTS) {
        for (coordinate in COORD) {
            frame = as.numeric(raw[-(1:3), 1])
            column = which(raw[2,] == part & raw[3,] == coordinate)
            value = as.numeric(raw[-(1:3), column])
            data[[i]] = data.frame(frame = frame, bodypart = part, 
                                   coordinate = coordinate, value = value)
            i = i + 1
        }
    }
    return(do.call(rbind, data))
}
likelihood_vaild = function(data) {
    data$valid = FALSE
    for (part in BODYPARTS) {
        likelihood_data = data[data$bodypart == part & data$coordinate == "likelihood",]
        index = data$bodypart == part
        match_index = match(data$frame[index], likelihood_data$frame)
        data$valid[index] = likelihood_data$value[match_index] >= LIKELIHOOD_THRESHOLD
    }
    return(data)
}
transform_recording = function(data, file_info) {
    head_x = data[data$bodypart == "head" & data$coordinate == "x", ]$value
    head_y = data[data$bodypart == "head" & data$coordinate == "y", ]$value
    head_valid = data[data$bodypart == "head" & data$coordinate == "x", ]$valid

    tail_x = data[data$bodypart == "tail" & data$coordinate == "x", ]$value
    tail_y = data[data$bodypart == "tail" & data$coordinate == "y", ]$value
    tail_valid = data[data$bodypart == "tail" & data$coordinate == "x", ]$valid

    dx = head_x - tail_x
    dy = head_y - tail_y
    body_length = sqrt(dx^2 + dy^2)
    body_valid = head_valid & tail_valid & is.finite(body_length) & body_length > 0
    body_reference = median(body_length[body_valid], na.rm = TRUE)
    center_x = (head_x + tail_x) / 2
    center_y = (head_y + tail_y) / 2
    theta = atan2(dy, dx)
    return_data = data
    for (leg in LEGS) {
        x_centered = data[data$bodypart == leg & data$coordinate == "x", ]$value - center_x
        y_centered = data[data$bodypart == leg & data$coordinate == "y", ]$value - center_y
        leg_valid = data[data$bodypart == leg & data$coordinate == "x", ]$valid

        eta_px = cos(theta) * x_centered + sin(theta) * y_centered
        xi_px = -sin(theta) * x_centered + cos(theta) * y_centered
        if (file_info$mode == "walk") {
            xi_px = -xi_px
        }
        
        eta = eta_px / body_reference
        xi = xi_px / body_reference
        valid = leg_valid & head_valid & tail_valid & is.finite(eta) & is.finite(xi)

        eta_data = data.frame(frame = data[data$bodypart == "head" & data$coordinate == "x", ]$frame,
                              bodypart = leg, coordinate = "eta", value = eta, valid = valid, stringsAsFactors = FALSE)
        xi_data = data.frame(frame = data[data$bodypart == "head" & data$coordinate == "x", ]$frame,
                             bodypart = leg, coordinate = "xi", value = xi, valid = valid, stringsAsFactors = FALSE)
        return_data = rbind(return_data, eta_data, xi_data)
    }
    return(return_data)
}

plot_body_centered = function(data, file_info) {
    axis_at = seq(-0.6, 0.6, by = 0.2)
    axis_labels = sprintf("%.1f", axis_at)

    pdf(file.path(PLOT_DIR, paste0(file_info$id, "_body_centered_trajectories.pdf")),
        width = 6, height = 6)
    par(tck = 0.015, las = 1, mar = c(4.5, 4.8, 1.5, 1))

    plot(NA, xlim = c(-0.6, 0.6), ylim = c(-0.6, 0.6), asp = 1,
         xlab = "Longitudinal eta (body lengths)",
         ylab = "Lateral xi (body lengths)", axes = FALSE)

    axis(1, at = axis_at, labels = axis_labels)
    axis(2, at = axis_at, labels = axis_labels, las = 1)
    box()
    abline(h = 0, v = 0, lty = 2)

    for (leg in LEGS) {
        eta = data[data$bodypart == leg & data$coordinate == "eta" & data$valid,]
        xi = data[data$bodypart == leg & data$coordinate == "xi" & data$valid,]
        points(eta$value, xi$value, pch = 16, cex = 0.25, col = LEG_COLOR[leg])
    }
    dev.off()
}

plot_eta_time_series = function(data, file_info) {
    axis_at = seq(-0.6, 0.6, by = 0.2)
    axis_labels = sprintf("%.1f", axis_at)
    x_range = range(data$frame / FPS, finite = TRUE)

    pdf(file.path(PLOT_DIR, paste0(file_info$id, "_eta_time_series.pdf")),
        width = 8, height = 5)
    par(tck = 0.015, las = 1, mar = c(4.5, 4.8, 1.5, 1))

    plot(NA, xlim = x_range, ylim = c(-0.6, 0.6),
         xlab = "Time (s)", ylab = "Longitudinal eta (body lengths)",
         axes = FALSE)

    axis(1)
    axis(2, at = axis_at, labels = axis_labels, las = 1)
    box()
    abline(h = 0, lty = 2)

    for (leg in LEGS) {
        eta = data[data$bodypart == leg & data$coordinate == "eta",]
        eta$value[!eta$valid] = NA
        lines(eta$frame / FPS, eta$value, col = LEG_COLOR[leg])
    }

    dev.off()
}

files = list.files(RAW_DIR, pattern = "\\.csv$", full.names = TRUE, ignore.case = TRUE)
for (path in files) {
    file_info = make_file_info(path)
    raw_data = read_dlc(path)
    valid_data = likelihood_vaild(raw_data)
    transformed_data = transform_recording(valid_data, file_info)
    write.csv(transformed_data, file.path(RECORDING_DIR, paste0(file_info$id, "_result.csv")),
              row.names = FALSE)
    plot_body_centered(transformed_data, file_info)
    plot_eta_time_series(transformed_data, file_info)
}
```
