## Supplemental Methods for "Data-driven frame-by-frame analysis of surface swimming and terrestrial walking in the coastal beetle *Phaleria atriceps*": 02_sinusoid_fitting.R

```
RAW_DIR = file.path("scratch_results", "coordinate_transform", "recordings")
OUTPUT_DIR = file.path("scratch_results", "sinusoid_fitting")
PLOT_DIR = file.path(OUTPUT_DIR, "plot")
DATA_DIR = file.path(OUTPUT_DIR, "recordings")

FREQUENCY_MIN = 0.5
FREQUENCY_MAX = 20
FREQUENCY_STEP = 0.02
FREQUENCY_GRID = seq(FREQUENCY_MIN, FREQUENCY_MAX, by = FREQUENCY_STEP)
FPS = 1000
LEGS = c("L1", "L2", "L3", "R1", "R2", "R3")
LEG_COLORS = setNames(hcl.colors(length(LEGS), "Dark 3"), LEGS)

dir.create(OUTPUT_DIR, recursive = TRUE, showWarnings = FALSE)
dir.create(PLOT_DIR, recursive = TRUE, showWarnings = FALSE)
dir.create(DATA_DIR, recursive = TRUE, showWarnings = FALSE)

read_data = function(path) {
    return(read.csv(path, header = TRUE, stringsAsFactors = FALSE,
                    check.names = FALSE, na.strings = c("", "NA", "NaN", "nan")))
}

fit_at_frequency = function(t, y, frequency, id, mode, leg) {
    X = cbind(1, sin(2*pi*frequency*t), cos(2*pi*frequency*t))
    fit = lm.fit(X, y)
    if (fit$rank < 3 || any(!is.finite(fit$coefficients))) {
        return(NULL)
    }
    fitted = as.numeric(X %*% fit$coefficients)
    residual = y - fitted

    sse = sum(residual^2)
    sst = sum((y - mean(y))^2)
    r2 = 1 - sse/sst

    return(data.frame(frequency = frequency, r2 = r2, offset = fit$coefficients[1],
                      sin_coefficient = fit$coefficients[2], cos_coefficient = fit$coefficients[3],
                      id = id, mode = mode, leg = leg))
}

files = list.files(RAW_DIR, pattern = "\\.csv$", full.names = TRUE, ignore.case = TRUE)
for (path in files) {
    data = read_data(path)
    file_name = tools::file_path_sans_ext(basename(path))
    id = sub("_result$", "", file_name)
    mode = sub("_.*$", "", id)
    all_profile = list()
    for (leg in LEGS) {
        leg_profile = list()
        eta = data[data$bodypart == leg & data$coordinate == "eta" & data$valid,]
        eta = eta[order(eta$frame), ]
        for (i in seq_along(FREQUENCY_GRID)) {
            freq = FREQUENCY_GRID[i]
            fit_result = fit_at_frequency(eta$frame / FPS, eta$value, freq, id, mode, leg)
            if (!is.null(fit_result)) {
                leg_profile[[i]] = fit_result
            }
        }
        leg_profile = do.call(rbind, leg_profile)
        all_profile[[leg]] = leg_profile
    }
    all_profile = do.call(rbind, all_profile)
    rownames(all_profile) = NULL
    write.csv(all_profile, file.path(DATA_DIR, paste0(id, "_result.csv")),
              row.names = FALSE)

}
```
