## Supplemental Methods for "Data-driven frame-by-frame analysis of surface swimming and terrestrial walking in the coastal beetle *Phaleria atriceps*": 02b_plot_representative_sinusoidal_fitting_gray.R

```
COORDINATE_DIR = file.path("scratch_results", "coordinate_transform", "recordings")
FITTING_DIR = file.path("scratch_results", "sinusoid_fitting", "recordings")
PERIOD_DIR = file.path("scratch_results", "selection_period", "recordings")
OUTPUT_DIR = file.path("scratch_results", "representative_sinusoidal_fitting")
PLOT_DIR = file.path(OUTPUT_DIR, "plot")
FPS = 1000
R2_THRESHOLD = 0.2
LEGS = c("L1", "L2", "L3", "R1", "R2", "R3")
LEG_COLORS = setNames(hcl.colors(length(LEGS), "Dark 3"), LEGS)
NONFOCAL_COLOR = "grey60"
NONFOCAL_ALPHA = 0.25
POSITION_X_MIN = -0.8
POSITION_X_MAX = 0.8
POSITION_X_AT = seq(POSITION_X_MIN, POSITION_X_MAX, by = 0.4)
POSITION_Y_MIN = -0.8
POSITION_Y_MAX = 0.8
POSITION_Y_AT = seq(POSITION_Y_MIN, POSITION_Y_MAX, by = 0.4)
R2_Y_MIN = 0
R2_Y_MAX = 1
R2_Y_AT = seq(R2_Y_MIN, R2_Y_MAX, by = 0.2)
REPRESENTATIVES = data.frame(
    panel_top = c("A", "B", "C", "D"),
    panel_middle = c("E", "F", "G", "H"),
    panel_bottom = c("I", "J", "K", "L"),
    id = c("swim_49", "swim_49", "swim_75", "walk_10"),
    leg = c("L3", "L1", "L3", "L2"),
    category = c("direct", "harmonic", "inconsistent", "direct"),
    stringsAsFactors = FALSE
)

dir.create(OUTPUT_DIR, recursive = TRUE, showWarnings = FALSE)
dir.create(PLOT_DIR, recursive = TRUE, showWarnings = FALSE)

read_data = function(path) {
    return(read.csv(path, header = TRUE, stringsAsFactors = FALSE,
                    check.names = FALSE, na.strings = c("", "NA", "NaN", "nan")))
}
get_mode_label = function(id) {
    mode = sub("_.*$", "", id)
    if (mode == "swim") {
        return("swimming")
    }
    if (mode == "walk") {
        return("walking")
    }
    return(mode)
}
make_body_panel_title = function(leg, mode_label) {
    return(paste0("Leg: ", leg, "\n", "Mode: ", mode_label))
}
make_time_series_title = function(selected, fitting_data, leg) {
    leg_fitting = fitting_data[fitting_data$leg == leg & is.finite(fitting_data$r2),]
    if (nrow(selected) > 0 && is.finite(selected$selected_frequency[1]) && is.finite(selected$selected_r2[1])) {
        frequency_text = paste0("Selected Frequency: ", sprintf("%.2f", selected$selected_frequency[1]))
        r2_value = selected$selected_r2[1]
    } else {
        frequency_text = "Selected Frequency: None"
        if (nrow(leg_fitting) > 0) {
            r2_value = max(leg_fitting$r2, na.rm = TRUE)
        } else {
            r2_value = NA_real_
        }
    }
    return(list(frequency_text = frequency_text, r2_value = r2_value))
}
draw_time_series_main = function(title_info) {
    if (is.finite(title_info$r2_value)) {
        title(main = bquote(atop(bold(.(title_info$frequency_text)),
                                 bold(R^2 * ": " *.(sprintf("%.2f", title_info$r2_value))))),
              cex.main = 2, family = "Arial")
    } else {
        title(main = bquote(atop(bold(.(title_info$frequency_text)),
                                 bold(R^2 * ": NA"))),
              cex.main = 1.2, family = "Arial")
    }
}
get_fit_row = function(fitting_data, leg, frequency) {
    fit_row = fitting_data[fitting_data$leg == leg & abs(fitting_data$frequency - frequency) < 1e-10,]
    if (nrow(fit_row) == 0) {
        return(NULL)
    }
    return(fit_row[1, ])
}
make_frequency_profile = function(fitting_data, leg) {
    data = fitting_data[fitting_data$leg == leg,]
    data = data[order(data$frequency),]
    rownames(data) = NULL
    return(data)
}
make_fitted_curve = function(time, fit_row, frequency) {
    fitted = fit_row$offset + fit_row$sin_coefficient * sin(2 * pi * frequency * time) +
             fit_row$cos_coefficient * cos(2 * pi * frequency * time)
    return(fitted)
}
draw_y_label = function(label) {
    mtext(label, side = 2, line = 5.0,
          cex = 1.5, las = 0, family = "Arial")
}
add_panel_label = function(label) {
    usr = par("usr")
    x = usr[1] - 0.13 * (usr[2] - usr[1])
    y = usr[4] + 0.12 * (usr[4] - usr[3])
    text(x, y, labels = label, adj = c(0, 1),
         cex = 5, font = 2, family = "Arial",
         xpd = NA)
}
draw_body_centered_panel = function(panel_info, coordinate_data) {
    mode_label = get_mode_label(panel_info$id)
    title_text = make_body_panel_title(panel_info$leg, mode_label)
    eta_all = coordinate_data[coordinate_data$bodypart %in% LEGS & 
                              coordinate_data$coordinate == "eta" & coordinate_data$valid &
                              is.finite(coordinate_data$value), c("frame", "bodypart", "value")]
    xi_all = coordinate_data[coordinate_data$bodypart %in% LEGS & coordinate_data$coordinate == "xi" &
                             coordinate_data$valid & is.finite(coordinate_data$value),
                             c("frame", "bodypart", "value")]
    names(eta_all)[3] = "longitudinal"
    names(xi_all)[3] = "lateral"
    merged = merge(eta_all, xi_all, by = c("frame", "bodypart"))
    merged = merged[order(merged$frame),]
    if (nrow(merged) == 0) {
        plot.new()
        title(main = title_text, cex.main = 2, font.main = 2, family = "Arial")
        add_panel_label(panel_info$panel_top)
        return(NULL)
    }
    plot(NA, NA, xlim = c(POSITION_X_MIN, POSITION_X_MAX), ylim = c(POSITION_Y_MIN, POSITION_Y_MAX),
        xlab = "Longitudinal position (body lengths)", ylab = "",
        main = title_text, asp = 1,
        family = "Arial", cex.lab = 2,
        cex.axis = 2, cex.main = 2,
        font.main = 2, xaxt = "n", yaxt = "n")
    axis(side = 1, at = POSITION_X_AT, labels = sprintf("%.1f", POSITION_X_AT), cex.axis = 2)
    axis(side = 2, at = POSITION_Y_AT, labels = sprintf("%.1f", POSITION_Y_AT), las = 1, cex.axis = 2)
    draw_y_label("Lateral position (body lengths)")
    abline(h = 0, v = 0, lty = 3, lwd = 1.5, col = "grey80")
    for (leg in LEGS) {
        leg_data = merged[merged$bodypart == leg,]
        if (nrow(leg_data) == 0) {
            next
        }
        if (leg == panel_info$leg) {
            points(leg_data$longitudinal, leg_data$lateral, pch = 16, cex = 1.0,
                   col = adjustcolor(LEG_COLORS[leg], alpha.f = 0.90))
        } else {
            points(leg_data$longitudinal, leg_data$lateral, pch = 16, cex = 0.65,
                col = adjustcolor(NONFOCAL_COLOR, alpha.f = NONFOCAL_ALPHA))
        }
    }
    add_panel_label(panel_info$panel_top)
    return(NULL)
}
draw_time_series_panel = function(panel_info, coordinate_data, fitting_data, selected_period) {
    selected = selected_period[selected_period$id == panel_info$id & selected_period$leg == panel_info$leg,]
    title_info = make_time_series_title(selected, fitting_data, panel_info$leg)
    profile_data = make_frequency_profile(fitting_data, panel_info$leg)
    eta = coordinate_data[coordinate_data$bodypart == panel_info$leg & coordinate_data$coordinate == "eta",]
    eta = eta[order(eta$frame),]
    time = eta$frame / FPS
    observed = eta$value
    observed[!eta$valid] = NA
    plot(time, observed, type = "n", xlab = "Time (s)", ylab = "", main = "",
         ylim = c(POSITION_Y_MIN, POSITION_Y_MAX), family = "Arial", cex.lab = 2, cex.axis = 2, yaxt = "n")
    draw_time_series_main(title_info)
    axis(side = 2, at = POSITION_Y_AT, labels = sprintf("%.1f", POSITION_Y_AT), las = 1, cex.axis = 2)
    draw_y_label("Longitudinal position (body lengths)")
    abline(h = 0, lty = 3, lwd = 1.5, col = "grey80")
    add_panel_label(panel_info$panel_middle)
    dominant_frequency = NA_real_
    dominant_fit = NULL
    if (nrow(profile_data) > 0) {
        max_row = profile_data[which.max(profile_data$r2),]
        dominant_frequency = max_row$frequency[1]
        dominant_fit = get_fit_row(fitting_data, panel_info$leg, dominant_frequency)
    }
    if (panel_info$category == "inconsistent") {
        if (!is.null(dominant_fit)) {
            dominant_curve = make_fitted_curve(time, dominant_fit, dominant_frequency)
            lines(time, dominant_curve, lwd = 3.5, lty = 1, col = "black")
        }
    } else {
        if (nrow(selected) > 0 && is.finite(selected$selected_frequency[1])) {
            selected_frequency = selected$selected_frequency[1]
            selected_fit = get_fit_row(fitting_data, panel_info$leg, selected_frequency)
            if (!is.null(selected_fit)) {
                selected_curve = make_fitted_curve(time, selected_fit, selected_frequency)
                if (panel_info$category == "harmonic" && !is.null(dominant_fit)) {
                    dominant_curve = make_fitted_curve(time, dominant_fit, dominant_frequency)
                    lines(time, dominant_curve, lwd = 2.5, lty = 2, col = "grey45")
                }
                lines(time, selected_curve, lwd = 3.5, lty = 1, col = "black")
            }
        }
    }
    points(time, observed, pch = 16, cex = 0.70,
           col = adjustcolor(LEG_COLORS[panel_info$leg],alpha.f = 0.55))
    return(NULL)
}
draw_frequency_profile_panel = function(panel_info, fitting_data, selected_period) {
    profile_data = make_frequency_profile(fitting_data, panel_info$leg)
    selected = selected_period[selected_period$id == panel_info$id & selected_period$leg == panel_info$leg,]
    if (nrow(profile_data) == 0) {
        plot.new()
        add_panel_label(panel_info$panel_bottom)
        return(NULL)
    }
    plot(profile_data$frequency, profile_data$r2, type = "l", lwd = 2.5, col = "grey35", 
         xlab = "Frequency (Hz)", ylab = "", main = "", ylim = c(R2_Y_MIN, R2_Y_MAX),
          family = "Arial", cex.lab = 2, cex.axis = 2, yaxt = "n")
    axis(side = 2, at = R2_Y_AT, labels = sprintf("%.1f", R2_Y_AT), las = 1, cex.axis = 2)
    draw_y_label(expression(R^2))
    abline(h = R2_THRESHOLD, lty = 3, lwd = 2, col = "grey60")
    max_row = profile_data[which.max(profile_data$r2),]
    points(max_row$frequency[1], max_row$r2[1], pch = 16, cex = 1.8, col = "black")
    if (nrow(selected) > 0 && is.finite(selected$reference_frequency[1])) {
        abline(v = selected$reference_frequency[1], lwd = 2.5, lty = 3, col = "black")
    }
    if (nrow(selected) > 0 && is.finite(selected$selected_frequency[1]) && is.finite(selected$selected_r2[1])) {
        points(selected$selected_frequency[1], selected$selected_r2[1], pch = 16, cex = 1.8, col = "red")
    }
    add_panel_label(panel_info$panel_bottom)
    return(NULL)
}
draw_all_panels = function(selected_period) {
    par(mfrow = c(3, 4), mar = c(5.5, 7.5, 6.0, 1.5), oma = c(1, 1, 1, 1), family = "Arial", las = 1, tck = 0.015)
    for (i in seq_len(nrow(REPRESENTATIVES))) {
        panel_info = REPRESENTATIVES[i,]
        coordinate_path = file.path(COORDINATE_DIR, paste0(panel_info$id, "_result.csv"))
        coordinate_data = read_data(coordinate_path)
        draw_body_centered_panel(panel_info, coordinate_data)
    }
    for (i in seq_len(nrow(REPRESENTATIVES))) {
        panel_info = REPRESENTATIVES[i,]
        coordinate_path = file.path(COORDINATE_DIR, paste0(panel_info$id, "_result.csv"))
        fitting_path = file.path(FITTING_DIR, paste0(panel_info$id, "_result.csv"))
        coordinate_data = read_data(coordinate_path)
        fitting_data = read_data(fitting_path)
        draw_time_series_panel(panel_info, coordinate_data, fitting_data, selected_period)
    }
    for (i in seq_len(nrow(REPRESENTATIVES))) {
        panel_info = REPRESENTATIVES[i,]
        fitting_path = file.path(FITTING_DIR, paste0(panel_info$id, "_result.csv"))
        fitting_data = read_data(fitting_path)
        draw_frequency_profile_panel(panel_info, fitting_data, selected_period)
    }
}
make_representative_figure = function() {
    selected_period = read_data(file.path(PERIOD_DIR, "selected_period.csv"))
    pdf_path = file.path(PLOT_DIR, "representative_sinusoidal_fitting.pdf")
    png_path = file.path(PLOT_DIR, "representative_sinusoidal_fitting.png")
    cairo_pdf(filename = pdf_path, width = 24, height = 18, family = "Arial")
    draw_all_panels(selected_period)
    dev.off()
    png(filename = png_path, width = 24, height = 18,
        units = "in", res = 300, type = "cairo", family = "Arial")
    draw_all_panels(selected_period)
    dev.off()
    return(c(pdf_path, png_path))
}
output_files = make_representative_figure()
print(output_files)
```
