## Supplemental Methods for "Data-driven frame-by-frame analysis of surface swimming and terrestrial walking in the coastal beetle *Phaleria atriceps*": 03_selection_period.R

```
RAW_DIR = file.path("scratch_results", "sinusoid_fitting", "recordings")
OUTPUT_DIR = file.path("scratch_results", "selection_period")
PLOT_DIR = file.path(OUTPUT_DIR, "plot")
DATA_DIR = file.path(OUTPUT_DIR, "recordings")

R2_THRESHOLD = 0.2
LEGS = c("L1", "L2", "L3", "R1", "R2", "R3")
LEG_COLORS = setNames(hcl.colors(length(LEGS), "Dark 3"), LEGS)

REFERENCE_TOLERANCE_RATIO = 1.30
PEAK_MATCH_TOLERANCE_RATIO = 1.30

dir.create(OUTPUT_DIR, recursive = TRUE, showWarnings = FALSE)
dir.create(PLOT_DIR, recursive = TRUE, showWarnings = FALSE)
dir.create(DATA_DIR, recursive = TRUE, showWarnings = FALSE)

read_data = function(path) {
    return(read.csv(path, header = TRUE, stringsAsFactors = FALSE,
                    check.names = FALSE, na.strings = c("", "NA", "NaN", "nan")))
}

synthesize_all_data = function(files) {
    all_profile = list()
    for (i in seq_along(files)) {
        all_profile[[i]] = read_data(files[i])
    }
    all_profile = do.call(rbind, all_profile)
    rownames(all_profile) = NULL
    return(all_profile)
}

make_candidate_peaks = function(all_data) {
    find_local_maxima = function(data) {
        data = data[order(data$frequency), ]
        if (nrow(data) < 3) {
            return(data.frame())
        }
        local_max = rep(FALSE, nrow(data))
        for (i in 2:(nrow(data) - 1)) {
            if (!is.finite(data$r2[i - 1]) || !is.finite(data$r2[i]) || !is.finite(data$r2[i + 1])) {
                next
            }
            if (data$r2[i] > data$r2[i - 1] && data$r2[i] > data$r2[i + 1]) {
                local_max[i] = TRUE
            }
        }
        return(data[local_max, ])
    }

    candidate_peaks = list()
    k = 1
    ids = unique(all_data$id)
    for (id in ids) {
        for (leg in LEGS) {
            data = all_data[all_data$id == id & all_data$leg == leg,]
            if (nrow(data) == 0) {
                next
            }
            peaks = find_local_maxima(data)
            if (nrow(peaks) == 0) {
                next
            }
            peaks = peaks[order(peaks$r2, decreasing = TRUE), ]
            peaks$peak_rank = seq_len(nrow(peaks))
            candidate_peaks[[k]] = peaks
            k = k + 1
        }
    }
    candidate_peaks = do.call(rbind, candidate_peaks)
    rownames(candidate_peaks) = NULL
    return(candidate_peaks)
}

make_max_r2_frequency = function(all_data) {
    max_r2_frequency = list()
    k = 1
    ids = unique(all_data$id)
    for (id in ids) {
        for (leg in LEGS) {
            data = all_data[all_data$id == id & all_data$leg == leg,]
            if (nrow(data) == 0) {
                next
            }
            data = data[is.finite(data$r2), ]
            if (nrow(data) == 0) {
                next
            }
            max_row = data[which.max(data$r2), ]
            max_row$fit_valid = max_row$r2 >= R2_THRESHOLD
            max_r2_frequency[[k]] = max_row
            k = k + 1
        }
    }
    max_r2_frequency = do.call(rbind, max_r2_frequency)
    rownames(max_r2_frequency) = NULL
    return(max_r2_frequency)
}

add_reference_frequency = function(candidate_peaks, max_r2_frequency) {
    candidate_peaks$reference_frequency = NA_real_
    ids = unique(candidate_peaks$id)
    for (id in ids) {
        mode = candidate_peaks$mode[candidate_peaks$id == id][1]
        for (leg in LEGS) {
            index = candidate_peaks$id == id & candidate_peaks$leg == leg
            if (!any(index)) {
                next
            }
            reference_data = max_r2_frequency[max_r2_frequency$id != id & max_r2_frequency$mode == mode &
                                              max_r2_frequency$leg == leg & max_r2_frequency$fit_valid,]
            if (nrow(reference_data) == 0) {
                next
            }
            reference_frequency = median(reference_data$frequency, na.rm = TRUE)
            candidate_peaks$reference_frequency[index] = reference_frequency
        }
    }
    return(candidate_peaks)
}

within_ratio = function(frequency, reference_frequency, tolerance_ratio) {
        if (!is.finite(frequency) || !is.finite(reference_frequency) ||
            frequency <= 0 || reference_frequency <= 0) {
            return(FALSE)
        }
        ratio = frequency / reference_frequency
        return(ratio >= 1 / tolerance_ratio & ratio <= tolerance_ratio)
}

add_reference_information = function(candidate_peaks) {    
    candidate_peaks$ratio_to_reference = candidate_peaks$frequency / candidate_peaks$reference_frequency
    candidate_peaks$reference_like = FALSE
    candidate_peaks$second_harmonic_like = FALSE
    candidate_peaks$half_frequency_like = FALSE

    for (i in seq_len(nrow(candidate_peaks))) {
        frequency = candidate_peaks$frequency[i]
        reference_frequency = candidate_peaks$reference_frequency[i]
        if (!is.finite(reference_frequency)) {
            next
        }
        candidate_peaks$reference_like[i] = within_ratio(frequency, reference_frequency, REFERENCE_TOLERANCE_RATIO)
        candidate_peaks$second_harmonic_like[i] = within_ratio(frequency / 2, reference_frequency, REFERENCE_TOLERANCE_RATIO)
        candidate_peaks$half_frequency_like[i] = within_ratio(frequency * 2, reference_frequency, REFERENCE_TOLERANCE_RATIO)
    }
    return(candidate_peaks)
}

select_period = function(candidate_peaks, max_r2_frequency) {
    selection = list()
    k = 1
    for (i in seq_len(nrow(max_r2_frequency))) {
        id = max_r2_frequency$id[i]
        mode = max_r2_frequency$mode[i]
        leg = max_r2_frequency$leg[i]

        selected_frequency = NA_real_
        selected_r2 = NA_real_
        selection_status = NA_character_

        peaks = candidate_peaks[candidate_peaks$id == id & candidate_peaks$leg == leg,]

        if (!max_r2_frequency$fit_valid[i]) {
            selection_status = "insufficient_fit_quality"
        } else if (nrow(peaks) == 0) {
            selection_status = "no_frequency_peak"
        } else {
            peaks = peaks[order(peaks$peak_rank), ]
            reference_frequency = peaks$reference_frequency[1]
            if (!is.finite(reference_frequency)) {
                selection_status = "reference_unavailable"
            } else {
                dominant = peaks[peaks$peak_rank == 1, ][1, ]
                if (dominant$reference_like) {
                    selected_frequency = dominant$frequency
                    selected_r2 = dominant$r2
                    selection_status = "direct_reference_consistent"
                } else if (dominant$second_harmonic_like) {
                    expected_fundamental = dominant$frequency / 2
                    fundamental_candidates = peaks[vapply(peaks$frequency,
                                                          function(frequency) {
                                                            within_ratio(frequency, expected_fundamental, PEAK_MATCH_TOLERANCE_RATIO)
                                                          }, logical(1)) & peaks$reference_like, ]
                    if (nrow(fundamental_candidates) == 0) {
                        selection_status = "possible_second_harmonic_without_fundamental_peak"
                    } else {
                        fundamental = fundamental_candidates[which.max(fundamental_candidates$r2), ]
                        selected_frequency = fundamental$frequency
                        selected_r2 = fundamental$r2
                        selection_status = "second_harmonic_corrected"
                    }
                } else {
                    selection_status = "reference_inconsistent"
                }
            }
        }
        selection[[k]] = data.frame(id = id, mode = mode, leg = leg,
                                    max_r2_frequency = max_r2_frequency$frequency[i],
                                    max_r2 = max_r2_frequency$r2[i],
                                    reference_frequency = if (nrow(peaks) > 0) peaks$reference_frequency[1] else NA_real_,
                                    selected_frequency = selected_frequency, selected_r2 = selected_r2,
                                    selection_status = selection_status, stringsAsFactors = FALSE)
        k = k + 1
    }
    selection = do.call(rbind, selection)
    rownames(selection) = NULL
    return(selection)
}

plot_frequency_profile = function(id, leg, all_data, candidate_peaks, max_r2_frequency,
                                  selected_period,plot_dir, top_n_peaks = 5) {
    profile = all_data[all_data$id == id & all_data$leg == leg,]
    if (nrow(profile) == 0) {
        return(NULL)
    }
    profile = profile[order(profile$frequency), ]
    peaks = candidate_peaks[candidate_peaks$id == id & candidate_peaks$leg == leg,]

    if (nrow(peaks) > 0) {
        peaks = peaks[order(peaks$r2, decreasing = TRUE), ]
        peaks_to_plot = peaks[seq_len(min(top_n_peaks, nrow(peaks))),]
        reference_frequency = peaks$reference_frequency[1]
    } else {
        peaks_to_plot = data.frame()
        reference_frequency = NA_real_
    }

    max_row = max_r2_frequency[max_r2_frequency$id == id & max_r2_frequency$leg == leg,]
    selected_row = selected_period[selected_period$id == id & selected_period$leg == leg,]

    selected_frequency = NA_real_
    selected_r2 = NA_real_

    if (nrow(selected_row) > 0 &&
        is.finite(selected_row$selected_frequency[1])) {
        selected_frequency = selected_row$selected_frequency[1]
        selected_r2 = selected_row$selected_r2[1]
    }

    mode = profile$mode[1]

    output_path = file.path(plot_dir, paste0(id, "_", leg, "_frequency_profile.png"))

    png(output_path, width = 1800, height = 1200, res = 200)

    par(tck = 0.015, las = 1)

    plot(profile$frequency, profile$r2, type = "l", lwd = 2, col = "black",
         xlab = "Frequency (Hz)", ylab = expression(R^2),
         main = paste0(id, " | ", mode, " | ", leg),
         ylim = range(c(0, profile$r2), finite = TRUE))

    if (nrow(peaks_to_plot) > 0) {
        points(peaks_to_plot$frequency, peaks_to_plot$r2, pch = 16, cex = 1.0, col = "black")
    }

    # if (nrow(max_row) > 0 && is.finite(max_row$frequency[1])) {
    #     abline(v = max_row$frequency[1], lty = 1, lwd = 1.5, col = "grey40")
    # }

    if (is.finite(reference_frequency)) {
        abline(v = reference_frequency, lty = 2, lwd = 1.5, col = "black")
    }

    if (is.finite(selected_frequency)) {
        abline(v = selected_frequency, lty = 2, lwd = 1.5, col = "red")
    }
    dev.off()
    return(output_path)
}

plot_all_frequency_profiles = function(all_data, candidate_peaks, max_r2_frequency,
                                       selected_period, plot_dir, top_n_peaks = 5) {
    output_files = list()
    k = 1
    ids = unique(all_data$id)
    for (id in ids) {
        for (leg in LEGS) {
            output_path = plot_frequency_profile(id = id, leg = leg, all_data = all_data,
                                                 candidate_peaks = candidate_peaks,
                                                 max_r2_frequency = max_r2_frequency,
                                                 selected_period = selected_period,
                                                 plot_dir = plot_dir, top_n_peaks = top_n_peaks)
            if (!is.null(output_path)) {
                output_files[[k]] = output_path
                k = k + 1
            }
        }
    }
    return(output_files)
}

files = list.files(RAW_DIR, pattern = "\\.csv$", full.names = TRUE, ignore.case = TRUE)
all_data = synthesize_all_data(files)
candidate_peaks = make_candidate_peaks(all_data)
max_r2_frequency = make_max_r2_frequency(all_data)
candidate_peaks_with_ref = add_reference_frequency(candidate_peaks, max_r2_frequency)
candidate_peaks_with_ref_info = add_reference_information(candidate_peaks_with_ref)
selected_period = select_period(candidate_peaks_with_ref_info, max_r2_frequency)
write.csv(selected_period, file.path(DATA_DIR, "selected_period.csv"), row.names = FALSE)
plot_files = plot_all_frequency_profiles(all_data = all_data, candidate_peaks = candidate_peaks_with_ref_info,
                                         max_r2_frequency = max_r2_frequency, selected_period = selected_period,
                                         plot_dir = PLOT_DIR, top_n_peaks = 5)
```
