## Supplemental Methods for "Data-driven frame-by-frame analysis of surface swimming and terrestrial walking in the coastal beetle *Phaleria atriceps*": 03b_plot_selected_period.R

```
COORDINATE_DIR = file.path("scratch_results", "coordinate_transform", "recordings")
FITTING_DIR = file.path("scratch_results", "sinusoid_fitting", "recordings")
PERIOD_DIR = file.path("scratch_results", "selection_period", "recordings")
OUTPUT_DIR = file.path("scratch_results","plot_selected_period" )
PLOT_DIR = file.path(OUTPUT_DIR, "plot")

FPS = 1000
LEGS = c("L1", "L2", "L3", "R1", "R2", "R3")
LEG_COLORS = setNames(hcl.colors(length(LEGS), "Dark 3"), LEGS)
AXIS_MAX = 0.6
AXIS_MIN = -0.6
AXIS_AT = seq(AXIS_MIN, AXIS_MAX, by = 0.2)

dir.create(OUTPUT_DIR, recursive = TRUE, showWarnings = FALSE)
dir.create(PLOT_DIR, recursive = TRUE, showWarnings = FALSE)

read_data = function(path) {
    return(read.csv(path, header = TRUE, stringsAsFactors = FALSE,
                    check.names = FALSE, na.strings = c("", "NA", "NaN", "nan")))
}

plot_selected_frequency = function(id, coordinate_data, fitting_data, selected_period) {
    output_path = file.path(PLOT_DIR, paste0(id, "_selected_frequency_fit.pdf"))
    eta_all = coordinate_data[coordinate_data$bodypart %in% LEGS & coordinate_data$coordinate == "eta",]
    if (nrow(eta_all) == 0) {
        return(NULL)
    }
    x_range = range(eta_all$frame / FPS, finite = TRUE)
    pdf(output_path, width = 10, height = 6)
    par(mar = c(4.5, 4.5, 2.5, 1), tck = 0.015, las = 1)
    plot(NA, xlim = x_range, ylim = c(AXIS_MIN, AXIS_MAX), xlab = "Time (s)", ylab = expression(eta), main = id)
    abline(h = 0, lty = 2, col = "grey70")
    for (leg in LEGS) {
        eta = coordinate_data[coordinate_data$bodypart == leg & coordinate_data$coordinate == "eta",]
        eta = eta[order(eta$frame), ]
        if (nrow(eta) == 0) {
            next
        }
        time = eta$frame / FPS
        observed = eta$value
        observed[!eta$valid] = NA
        raw_color = adjustcolor(LEG_COLORS[leg], alpha.f = 0.30)
        lines(time, observed, lwd = 1, col = raw_color)
        points(time[eta$valid], eta$value[eta$valid], pch = 16, cex = 0.25, col = raw_color)
        selected = selected_period[selected_period$id == id & selected_period$leg == leg,]
        if (nrow(selected) == 0 || !is.finite(selected$selected_frequency[1])) {
            next
        }
        selected_frequency = selected$selected_frequency[1]
        fit_row = fitting_data[fitting_data$leg == leg & abs(fitting_data$frequency - selected_frequency) < 1e-10,]
        if (nrow(fit_row) == 0) {
            next
        }
        fit_row = fit_row[1, ]
        fitted = fit_row$offset + fit_row$sin_coefficient * sin(2*pi*selected_frequency*time) +
                                  fit_row$cos_coefficient * cos(2*pi*selected_frequency*time)
        lines(time, fitted, lwd = 2, col = LEG_COLORS[leg])
    }
    dev.off()
    return(output_path)
}


selected_period = read_data(paste0(PERIOD_DIR, "/selected_period.csv"))
coordinate_files = list.files(COORDINATE_DIR, pattern = "\\.csv$", full.names = TRUE, ignore.case = TRUE)

for (coordinate_path in coordinate_files) {
    file_name = tools::file_path_sans_ext(basename(coordinate_path))
    id = sub("_result$", "", file_name)
    fitting_path = file.path(FITTING_DIR, paste0(id, "_result.csv"))
    if (!file.exists(fitting_path)) {
        next
    }
    coordinate_data = read_data(coordinate_path)
    fitting_data = read_data(fitting_path)
    plot_selected_frequency(id = id, coordinate_data = coordinate_data,
                            fitting_data = fitting_data, selected_period = selected_period)
}
```
