## Supplemental Methods for "Data-driven frame-by-frame analysis of surface swimming and terrestrial walking in the coastal beetle *Phaleria atriceps*": 04_period_statistics.R

```
PERIOD_DIR = file.path("scratch_results", "selection_period", "recordings")
OUTPUT_DIR = file.path("scratch_results", "period_statistics")
DATA_DIR = file.path(OUTPUT_DIR, "recordings")
LEGS = c("L1", "L2", "L3", "R1", "R2", "R3")
MODES = c("swim", "walk")

dir.create(OUTPUT_DIR, recursive = TRUE, showWarnings = FALSE)
dir.create(DATA_DIR, recursive = TRUE, showWarnings = FALSE)

read_data = function(path) {
    return(read.csv(path, header = TRUE, stringsAsFactors = FALSE,
                    check.names = FALSE, na.strings = c("", "NA", "NaN", "nan")))
}

make_period_data = function(selected_period) {
    period_data = selected_period[is.finite(selected_period$selected_frequency) & 
                                  selected_period$selected_frequency > 0,]
    period_data$period = 1 / period_data$selected_frequency
    period_data$log_period = log(period_data$period)
    period_data$mode = factor(period_data$mode, levels = MODES)
    period_data$leg = factor(period_data$leg, levels = LEGS)
    period_data$id = factor(period_data$id)
    rownames(period_data) = NULL
    return(period_data)
}

fit_period_model = function(period_data) {
    model = nlme::lme(fixed = log_period ~ mode * leg, random = ~ 1 | id, weights = nlme::varIdent(form = ~ 1 | mode),
                      data = period_data, method = "REML", na.action = na.omit)
    return(model)
}


make_period_contrasts = function(model) {
    emmeans_result = emmeans::emmeans(model, ~ mode | leg)
    contrast_result = emmeans::contrast(emmeans_result, method = list("walk - swim" = c(-1, 1)), adjust = "none")
    contrast_result = as.data.frame(summary(contrast_result, infer = c(TRUE, TRUE), adjust = "none"))
    contrast_result$p_value_raw = contrast_result$p.value
    contrast_result$p_value_holm = p.adjust(contrast_result$p_value_raw, method = "holm")
    contrast_result$p.value = NULL
    contrast_result$period_ratio = exp(contrast_result$estimate)
    contrast_result$period_ratio_lower = exp(contrast_result$lower.CL)
    contrast_result$period_ratio_upper = exp(contrast_result$upper.CL)
    return(contrast_result)
}

make_period_summary = function(period_data) {
    summary_data = list()
    k = 1
    for (mode in MODES) {
        for (leg in LEGS) {
            data = period_data[period_data$mode == mode & period_data$leg == leg,]
            if (nrow(data) == 0) {
                next
            }
            summary_data[[k]] = data.frame(mode = mode, leg = leg, n = nrow(data),
                                           mean_period = mean(data$period), sd_period = sd(data$period),
                                           median_period = median(data$period),
                                           mean_frequency = mean(data$selected_frequency),
                                           sd_frequency = sd(data$selected_frequency),
                                           stringsAsFactors = FALSE)
            k = k + 1
        }
    }
    summary_data = do.call(rbind, summary_data)
    rownames(summary_data) = NULL
    return(summary_data)
}


selected_period = read_data(file.path(PERIOD_DIR, "selected_period.csv"))
period_data = make_period_data(selected_period)
period_model = fit_period_model(period_data)
period_anova = as.data.frame(anova(period_model))
period_contrasts = make_period_contrasts(period_model)
period_summary = make_period_summary(period_data)
write.csv(period_data, file.path(DATA_DIR, "period_data.csv"), row.names = FALSE)
write.csv(period_anova, file.path(DATA_DIR, "period_anova.csv"), row.names = TRUE)
write.csv(period_contrasts, file.path(DATA_DIR, "period_swim_walk_contrasts.csv"), row.names = FALSE)
write.csv(period_summary, file.path(DATA_DIR, "period_summary.csv"), row.names = FALSE)

print(summary(period_model))
print(anova(period_model))
print(period_contrasts)
print(period_summary)
```
