## Supplemental Methods for "Data-driven frame-by-frame analysis of surface swimming and terrestrial walking in the coastal beetle *Phaleria atriceps*": 04b_plot_period_statistics.R

```
PERIOD_DIR = file.path("scratch_results", "period_statistics", "recordings")
OUTPUT_DIR = file.path("scratch_results", "period_statistics_plot")
PLOT_DIR = file.path(OUTPUT_DIR, "plot")
LEGS = c("L1", "L2", "L3", "R1", "R2", "R3")
MODES = c("swim", "walk")
MODE_COLORS = c(swim = "#1F77B4", walk = "#D62728")

dir.create(OUTPUT_DIR, recursive = TRUE, showWarnings = FALSE)
dir.create(PLOT_DIR, recursive = TRUE, showWarnings = FALSE)
read_data = function(path) {
    return(read.csv(path, header = TRUE, stringsAsFactors = FALSE,
                    check.names = FALSE, na.strings = c("", "NA", "NaN", "nan")))
}
fit_period_model = function(period_data) {
    model = nlme::lme(fixed = log_period ~ mode * leg, random = ~ 1 | id,
                      weights = nlme::varIdent(form = ~ 1 | mode),
                      data = period_data, method = "REML", na.action = na.omit)
    return(model)
}
make_emmeans_period = function(model) {
    emmeans_result = emmeans::emmeans(model, ~ mode | leg)
    emmeans_result = as.data.frame(summary(emmeans_result, infer = c(TRUE, TRUE)))
    emmeans_result$period = exp(emmeans_result$emmean)
    emmeans_result$period_lower = exp(emmeans_result$lower.CL)
    emmeans_result$period_upper = exp(emmeans_result$upper.CL)
    return(emmeans_result)
}
p_to_star = function(p) {
    if (!is.finite(p)) {return("")}
    if (p < 0.001) {return("***")}
    if (p < 0.01) {return("**")}
    if (p < 0.05) {return("*")}
    return("ns")
}
plot_period_one_panel = function(period_data, period_emmeans, period_contrasts) {
    output_path = file.path(PLOT_DIR, "period_by_leg_one_panel.pdf")
    period_data$leg = factor(period_data$leg, levels = LEGS)
    period_data$mode = factor(period_data$mode, levels = MODES)
    period_emmeans$leg = factor(period_emmeans$leg, levels = LEGS)
    period_emmeans$mode = factor(period_emmeans$mode, levels = MODES)
    period_contrasts$leg = factor(period_contrasts$leg, levels = LEGS)
    x_center = seq_along(LEGS)
    names(x_center) = LEGS
    offset = 0.18
    x_swim = x_center - offset
    x_walk = x_center + offset
    pdf(output_path, width = 10, height = 5.8)
    par(mar = c(5, 4.5, 2.5, 1), las = 1, tck = 0.015)
    plot(NA, NA, xlim = c(0.5, length(LEGS) + 0.5), ylim = c(0, 0.2),
        xaxt = "n", yaxt = "n", xlab = "Leg", ylab = "Period (s)", bty = "n")

    axis(1, at = x_center, labels = LEGS, lwd = 1.5)
    axis(2, at = seq(0, 0.2, by = 0.05), labels = sprintf("%.2f", seq(0, 0.2, by = 0.05)), las = 1, lwd = 1.5)
    for (leg in LEGS) {
        xc = x_center[leg]
        xs = x_swim[leg]
        xw = x_walk[leg]
        data_leg = period_data[period_data$leg == leg,]
        em_leg = period_emmeans[period_emmeans$leg == leg,]
        contrast_leg = period_contrasts[period_contrasts$leg == leg,]
        data_swim = data_leg[data_leg$mode == "swim",]
        data_walk = data_leg[data_leg$mode == "walk",]
        if (nrow(data_swim) > 0) {
            points(jitter(rep(xs, nrow(data_swim)), amount = 0.04),
                   data_swim$period, pch = 16, cex = 0.8,
                   col = adjustcolor(MODE_COLORS["swim"], alpha.f = 0.45))
        }
        if (nrow(data_walk) > 0) {
            points(jitter(rep(xw, nrow(data_walk)), amount = 0.04),
                   data_walk$period, pch = 16, cex = 0.8,
                   col = adjustcolor(MODE_COLORS["walk"], alpha.f = 0.45))
        }
        em_swim = em_leg[em_leg$mode == "swim",]
        em_walk = em_leg[em_leg$mode == "walk",]
        if (nrow(em_swim) == 1) {
            segments(xs, em_swim$period_lower, xs, em_swim$period_upper, lwd = 2, col = MODE_COLORS["swim"])
            segments(xs - 0.05, em_swim$period_lower, xs + 0.05, em_swim$period_lower, lwd = 2, col = MODE_COLORS["swim"])
            segments(xs - 0.05, em_swim$period_upper, xs + 0.05, em_swim$period_upper, lwd = 2, col = MODE_COLORS["swim"])
            points(xs, em_swim$period, pch = 18, cex = 1.4, col = MODE_COLORS["swim"])
        }
        if (nrow(em_walk) == 1) {
            segments(xw, em_walk$period_lower, xw, em_walk$period_upper, lwd = 2, col = MODE_COLORS["walk"])
            segments(xw - 0.05, em_walk$period_lower, xw + 0.05, em_walk$period_lower, lwd = 2, col = MODE_COLORS["walk"])
            segments(xw - 0.05, em_walk$period_upper,xw + 0.05, em_walk$period_upper,lwd = 2, col =MODE_COLORS["walk"])
            points(xw, em_walk$period, pch = 18, cex = 1.4, col =MODE_COLORS["walk"])
        }
        y_candidates =c(data_leg$period, em_leg$period_upper)
        y_sig =min(0.192, max(y_candidates, na.rm = TRUE) + 0.012)
        segments(xs, y_sig, xw, y_sig, lwd = 1.3, col = "black")
        segments(xs, y_sig - 0.004, xs, y_sig, lwd = 1.3, col = "black")
        segments(xw, y_sig - 0.004, xw, y_sig, lwd = 1.3, col = "black")
        if (nrow(contrast_leg) == 1) {
            if ("p_value_holm" %in% colnames(contrast_leg)) {
                p_value = contrast_leg$p_value_holm
            } else if ("p.value" %in% colnames(contrast_leg)) {
                p_value =contrast_leg$p.value
            } else {
                p_value =NA_real_
            }
            text(xc, y_sig + 0.005, labels = p_to_star(p_value), cex = 1.2)
        }
    }
    dev.off()
    return(output_path)
}

period_data =read_data(file.path(PERIOD_DIR,"period_data.csv"))
period_contrasts =read_data(file.path(PERIOD_DIR,"period_swim_walk_contrasts.csv"))
period_data$mode =factor(period_data$mode,levels = MODES)
period_data$leg =factor(period_data$leg,levels = LEGS)
period_model =fit_period_model(period_data)
period_emmeans =make_emmeans_period(period_model)
plot_period_one_panel(period_data,period_emmeans,period_contrasts)
```
