## Supplemental Methods for "Data-driven frame-by-frame analysis of surface swimming and terrestrial walking in the coastal beetle *Phaleria atriceps*": 05_phase_detect.R

```
COORDINATE_DIR = file.path("scratch_results", "coordinate_transform", "recordings")
FITTING_DIR = file.path("scratch_results", "sinusoid_fitting", "recordings")
PERIOD_DIR = file.path("scratch_results", "selection_period", "recordings")

OUTPUT_DIR = file.path("scratch_results", "phase_analysis")
DATA_DIR = file.path(OUTPUT_DIR, "recordings")

FPS = 1000
R2_THRESHOLD = 0.2

LEGS = c("L1", "L2", "L3", "R1", "R2", "R3")
TRIPOD_A = c("L1", "R2", "L3")
TRIPOD_B = c("R1", "L2", "R3")

dir.create(OUTPUT_DIR, recursive = TRUE, showWarnings = FALSE)
dir.create(DATA_DIR, recursive = TRUE, showWarnings = FALSE)

read_data = function(path) {
    return(read.csv(path, header = TRUE, stringsAsFactors = FALSE,
                    check.names = FALSE, na.strings = c("", "NA", "NaN", "nan")))
}
wrap_pi = function(angle) {
    return(atan2(sin(angle), cos(angle)))
}
rad_to_deg = function(angle) {
    return(180 * angle / pi)
}

get_selected_fit = function(fitting_data, leg, selected_frequency) {
    fit_row = fitting_data[fitting_data$leg == leg &
                           abs(fitting_data$frequency - selected_frequency) < 1e-10,]
    if (nrow(fit_row) == 0) {
        return(NULL)
    }
    return(fit_row[1,])
}

make_phase_parameters = function(selected_period) {
    phase_parameters = list()
    k = 1
    ids = unique(selected_period$id)
    for (id in ids) {
        coordinate_path = file.path(COORDINATE_DIR, paste0(id, "_result.csv"))
        fitting_path = file.path(FITTING_DIR, paste0(id, "_result.csv"))
        if (!file.exists(coordinate_path) || !file.exists(fitting_path)) {
            next
        }
        coordinate_data = read_data(coordinate_path)
        fitting_data = read_data(fitting_path)
        for (leg in LEGS) {
            selected = selected_period[selected_period$id == id &
                                       selected_period$leg == leg,]
            if (nrow(selected) == 0) {
                next
            }
            selected_frequency = selected$selected_frequency[1]
            if (!is.finite(selected_frequency)) {
                next
            }
            fit_row = get_selected_fit(fitting_data, leg, selected_frequency)
            if (is.null(fit_row)) {
                next
            }
            eta = coordinate_data[coordinate_data$bodypart == leg &
                                  coordinate_data$coordinate == "eta" &
                                  coordinate_data$valid &
                                  is.finite(coordinate_data$frame) &
                                  is.finite(coordinate_data$value),]
            if (nrow(eta) == 0) {
                next
            }
            eta = eta[order(eta$frame),]
            support_start = min(eta$frame) / FPS
            support_end = max(eta$frame) / FPS
            phase_offset = atan2(fit_row$cos_coefficient, fit_row$sin_coefficient)
            amplitude = sqrt(fit_row$sin_coefficient^2 + fit_row$cos_coefficient^2)
            phase_parameters[[k]] = data.frame(id = id, mode = selected$mode[1],
                                               leg = leg, frequency = selected_frequency,
                                               offset = fit_row$offset,
                                               sin_coefficient = fit_row$sin_coefficient,
                                               cos_coefficient = fit_row$cos_coefficient,
                                               phase_offset_rad = phase_offset,
                                               r2 = fit_row$r2,
                                               support_start = support_start,
                                               support_end = support_end,
                                               phase_valid = is.finite(fit_row$r2) & fit_row$r2 >= R2_THRESHOLD,
                                               stringsAsFactors = FALSE)
            k = k + 1
        }
    }
    if (length(phase_parameters) == 0) {
        return(data.frame())
    }
    phase_parameters = do.call(rbind, phase_parameters)
    rownames(phase_parameters) = NULL
    return(phase_parameters)
}


pair_relation = function(leg1, leg2) {
    if ((leg1 %in% TRIPOD_A && leg2 %in% TRIPOD_A) ||
        (leg1 %in% TRIPOD_B && leg2 %in% TRIPOD_B)) {
        return("same_tripod")
    }
    return("opposite_tripod")
}


ideal_phase = function(leg1, leg2) {
    if (pair_relation(leg1, leg2) == "same_tripod") {
        return(0)
    }
    return(pi)
}


make_relative_phase = function(phase_parameters) {
    relative_phase = list()
    k = 1
    leg_pairs = combn(LEGS, 2, simplify = FALSE)
    ids = unique(phase_parameters$id)
    for (id in ids) {
        data = phase_parameters[phase_parameters$id == id & phase_parameters$phase_valid,]
        for (pair in leg_pairs) {
            leg1 = pair[1]
            leg2 = pair[2]
            phase1 = data[data$leg == leg1,]
            phase2 = data[data$leg == leg2,]
            if (nrow(phase1) != 1 || nrow(phase2) != 1) {
                next
            }
            overlap_start = max(phase1$support_start, phase2$support_start)
            overlap_end = min(phase1$support_end, phase2$support_end)
            if (!is.finite(overlap_start) || !is.finite(overlap_end) || overlap_end <= overlap_start) {
                next
            }
            midpoint = (overlap_start + overlap_end) / 2
            theta1 = 2*pi*phase1$frequency*midpoint + phase1$phase_offset_rad
            theta2 = 2*pi*phase2$frequency*midpoint + phase2$phase_offset_rad
            delta_phase = wrap_pi(theta2 - theta1)
            ideal = ideal_phase(leg1, leg2)
            tripod_error = wrap_pi(delta_phase - ideal)
            relative_phase[[k]] = data.frame(id = id, mode = phase1$mode[1],
                                             pair = paste0(leg1, "-", leg2),
                                             leg1 = leg1, leg2 = leg2,
                                             tripod_relation = pair_relation(leg1, leg2),
                                             frequency_leg1 = phase1$frequency,
                                             frequency_leg2 = phase2$frequency,
                                             phase_offset_leg1_rad = phase1$phase_offset_rad,
                                             phase_offset_leg2_rad = phase2$phase_offset_rad,
                                             overlap_start = overlap_start, overlap_end = overlap_end, midpoint = midpoint,
                                             relative_phase_rad = delta_phase,
                                             ideal_phase_rad = ideal,
                                             tripod_error_rad = tripod_error,
                                             stringsAsFactors = FALSE)
            k = k + 1
        }
    }
    if (length(relative_phase) == 0) {
        return(data.frame())
    }
    relative_phase = do.call(rbind, relative_phase)
    rownames(relative_phase) = NULL
    return(relative_phase)
}


selected_period = read_data(file.path(PERIOD_DIR, "selected_period.csv"))
phase_parameters = make_phase_parameters(selected_period)
relative_phase = make_relative_phase(phase_parameters)
write.csv(phase_parameters, file.path(DATA_DIR, "phase_parameters.csv"), row.names = FALSE)
write.csv(relative_phase, file.path(DATA_DIR, "relative_phase.csv"), row.names = FALSE)


print(head(phase_parameters, 20))
print(head(relative_phase, 20))
```
