## Supplemental Methods for "Data-driven frame-by-frame analysis of surface swimming and terrestrial walking in the coastal beetle *Phaleria atriceps*": 06_phase_swim_walk_statistics.R

```
PHASE_DIR = file.path("scratch_results", "phase_analysis", "recordings")

OUTPUT_DIR = file.path("scratch_results", "phase_swim_walk_statistics")
DATA_DIR = file.path(OUTPUT_DIR, "recordings")

LEGS = c("L1", "L2", "L3", "R1", "R2", "R3")
MODES = c("swim", "walk")

TOLERANCE = 1e-12

dir.create(OUTPUT_DIR, recursive = TRUE, showWarnings = FALSE)
dir.create(DATA_DIR, recursive = TRUE, showWarnings = FALSE)


read_data = function(path) {
    return(read.csv(path, header = TRUE, stringsAsFactors = FALSE,
                    check.names = FALSE, na.strings = c("", "NA", "NaN", "nan")))
}
wrap_pi = function(angle) {
    return(atan2(sin(angle), cos(angle)))
}
rad_to_deg = function(angle) {
    return(180 * angle / pi)
}
rad_to_deg_360 = function(angle) {
    degree = (180 * angle / pi) %% 360
    degree[degree < 0] = degree[degree < 0] + 360
    return(degree)
}
circular_mean = function(angle) {
    angle = angle[is.finite(angle)]
    if (length(angle) == 0) {
        return(NA_real_)
    }
    return(atan2(mean(sin(angle)), mean(cos(angle))))
}
resultant_length = function(angle) {
    angle = angle[is.finite(angle)]
    if (length(angle) == 0) {
        return(NA_real_)
    }
    mean_sin = mean(sin(angle))
    mean_cos = mean(cos(angle))
    return(sqrt(mean_sin^2 + mean_cos^2))
}
make_group_matrix = function(n, n_swim) {
    combinations = combn(seq_len(n), n_swim)
    n_permutations = ncol(combinations)
    group_matrix = matrix(0, nrow = n_permutations, ncol = n)
    for (j in seq_len(n_swim)) {
        group_matrix[cbind(seq_len(n_permutations), combinations[j,])] = 1
    }
    return(group_matrix)
}


exact_phase_test = function(angle, mode) {
    valid = is.finite(angle) & mode %in% MODES
    angle = angle[valid]
    mode = as.character(mode[valid])
    n = length(angle)
    n_swim = sum(mode == "swim")
    n_walk = sum(mode == "walk")
    if (n_swim < 2 || n_walk < 2) {
        return(NULL)
    }
    mu_swim = circular_mean(angle[mode == "swim"])
    mu_walk = circular_mean(angle[mode == "walk"])
    phase_difference = wrap_pi(mu_swim - mu_walk)
    statistic_observed = 1 - cos(phase_difference)
    group_matrix = make_group_matrix(n, n_swim)
    sin_angle = sin(angle)
    cos_angle = cos(angle)
    sum_sin_swim = drop(group_matrix %*% sin_angle)
    sum_cos_swim = drop(group_matrix %*% cos_angle)
    total_sin = sum(sin_angle)
    total_cos = sum(cos_angle)
    sum_sin_walk = total_sin - sum_sin_swim
    sum_cos_walk = total_cos - sum_cos_swim
    mu_swim_permutation = atan2(sum_sin_swim, sum_cos_swim)
    mu_walk_permutation = atan2(sum_sin_walk, sum_cos_walk)
    statistic_permutation = 1 - cos(mu_swim_permutation - mu_walk_permutation)
    p_exact = mean(statistic_permutation >= statistic_observed - TOLERANCE)
    return(data.frame(n_swim = n_swim, n_walk = n_walk,
                      swim_mean_phase_rad = mu_swim, walk_mean_phase_rad = mu_walk,
                      swim_R = resultant_length(angle[mode == "swim"]),
                      walk_R = resultant_length(angle[mode == "walk"]),
                      swim_minus_walk_phase_rad = phase_difference,
                      absolute_phase_difference_rad = abs(phase_difference),
                      phase_difference_statistic = statistic_observed,
                      p_exact = p_exact, n_exact_permutations = nrow(group_matrix),
                      stringsAsFactors = FALSE))
}

make_phase_comparisons = function(relative_phase) {
    leg_pairs = combn(LEGS, 2, simplify = FALSE)
    result = list()
    k = 1
    for (pair in leg_pairs) {
        pair_name = paste0(pair[1], "-", pair[2])
        data = relative_phase[relative_phase$pair == pair_name,]
        if (nrow(data) == 0) {
            next
        }
        test = exact_phase_test(data$relative_phase_rad, data$mode)
        if (is.null(test)) {
            next
        }
        test$pair = pair_name
        test$leg1 = pair[1]
        test$leg2 = pair[2]
        result[[k]] = test
        k = k + 1
    }
    if (length(result) == 0) {
        return(data.frame())
    }
    result = do.call(rbind, result)
    rownames(result) = NULL
    result = result[, c("pair", "leg1", "leg2", setdiff(names(result), c("pair", "leg1", "leg2")))]
    result$swim_mean_phase_deg = rad_to_deg_360(result$swim_mean_phase_rad)
    result$walk_mean_phase_deg = rad_to_deg_360(result$walk_mean_phase_rad)
    result$swim_minus_walk_phase_deg = rad_to_deg(result$swim_minus_walk_phase_rad)
    result$absolute_phase_difference_deg = rad_to_deg(result$absolute_phase_difference_rad)
    result$p_holm = p.adjust(result$p_exact, method = "holm")
    result$significant = result$p_holm < 0.05
    return(result)
}

relative_phase = read_data(
    file.path(PHASE_DIR, "relative_phase.csv")
)

if (any(duplicated(relative_phase[, c("id", "pair")]))) {
    stop("Duplicated id x pair observations were found.")
}

phase_comparisons = make_phase_comparisons(relative_phase)
write.csv(phase_comparisons, file.path(DATA_DIR, "swim_walk_phase_comparisons.csv"), row.names = FALSE)

print(phase_comparisons)
```
