## Supplemental Methods for "Data-driven frame-by-frame analysis of surface swimming and terrestrial walking in the coastal beetle *Phaleria atriceps*": 06b_plot_phase_swim_vs_walk.R

```
RELATIVE_PHASE_DIR = file.path("scratch_results", "phase_analysis", "recordings")
PHASE_STAT_DIR = file.path("scratch_results", "phase_swim_walk_statistics", "recordings")
OUTPUT_DIR = file.path("scratch_results", "plot_phase_swim_vs_walk")
PLOT_DIR = file.path(OUTPUT_DIR, "plot")

LEGS = c("L1", "L2", "L3", "R1", "R2", "R3")
MODE_COLORS = c(swim = "#1F77B4", walk = "#D62728")

POINT_RADIUS = c(swim = 1.00, walk = 0.82)
ARROW_SCALE = 0.72

ANGLE_LABEL_RADIUS = 1.25
ANGLE_LABEL_CEX = 1.4
MAIN_TEXT_CEX = 1.35
PANEL_LABEL_CEX = 3.0
SIGNIFICANCE_CEX = 1.5
SIGNIFICANCE_OFFSET = 0.15
PLOT_LIMIT = 1.35

dir.create(OUTPUT_DIR, recursive = TRUE, showWarnings = FALSE)
dir.create(PLOT_DIR, recursive = TRUE, showWarnings = FALSE)

read_data = function(path) {
    return(read.csv(path, header = TRUE, stringsAsFactors = FALSE,
                    check.names = FALSE, na.strings = c("", "NA", "NaN", "nan")))
}
p_to_star = function(p) {
    if (!is.finite(p)) {return("")}
    if (p < 0.001) {return("***")}
    if (p < 0.01) {return("**")}
    if (p < 0.05) {return("*")}
    return("")
}
angle_to_xy = function(angle, radius = 1) {
    x = radius * cos(angle)
    y = radius * sin(angle)
    return(data.frame(x = x, y = y))
}
draw_phase_circle = function(pair_name, relative_phase, comparison_row, panel_label) {
    plot(NA, NA,
         xlim = c(-PLOT_LIMIT, PLOT_LIMIT),
         ylim = c(-PLOT_LIMIT, PLOT_LIMIT),
         asp = 1,
         xaxt = "n", yaxt = "n",
         xlab = "", ylab = "",
         bty = "n")

    theta = seq(0, 2*pi, length.out = 361)
    lines(cos(theta), sin(theta), lwd = 1.1, col = "black")

    segments(-1, 0, 1, 0, lwd = 0.8, col = "grey75")
    segments(0, -1, 0, 1, lwd = 0.8, col = "grey75")

    text(ANGLE_LABEL_RADIUS, 0, "0°", cex = ANGLE_LABEL_CEX)
    text(0, ANGLE_LABEL_RADIUS, "90°", cex = ANGLE_LABEL_CEX)
    text(-ANGLE_LABEL_RADIUS, 0, "180°", cex = ANGLE_LABEL_CEX)
    text(0, -ANGLE_LABEL_RADIUS, "270°", cex = ANGLE_LABEL_CEX)

    pair_data = relative_phase[
        relative_phase$pair == pair_name &
        is.finite(relative_phase$relative_phase_rad),
    ]

    star = ""
    if (nrow(comparison_row) == 1) {
        star = p_to_star(comparison_row$p_holm[1])
    }

    title(main = pair_name, line = 0.2, cex.main = MAIN_TEXT_CEX)

    usr = par("usr")
    x_range = usr[2] - usr[1]
    y_range = usr[4] - usr[3]

    text(usr[1] + 0.015*x_range,
         usr[4] - 0.015*y_range,
         labels = panel_label,
         adj = c(0, 1),
         cex = PANEL_LABEL_CEX,
         font = 2,
         xpd = NA)

    arrow_endpoints = list()

    for (mode in c("swim", "walk")) {
        data_mode = pair_data[pair_data$mode == mode, ]

        if (nrow(data_mode) == 0) {
            next
        }

        point_xy = angle_to_xy(data_mode$relative_phase_rad, POINT_RADIUS[mode])

        points(point_xy$x, point_xy$y,
               pch = 16,
               cex = 0.85,
               col = adjustcolor(MODE_COLORS[mode], alpha.f = 0.65))

        if (nrow(comparison_row) == 1) {
            mean_angle = comparison_row[[paste0(mode, "_mean_phase_rad")]][1]
            mean_r = comparison_row[[paste0(mode, "_R")]][1]

            if (is.finite(mean_angle) && is.finite(mean_r)) {
                arrow_length = ARROW_SCALE * mean_r
                arrow_xy = angle_to_xy(mean_angle, arrow_length)

                arrows(0, 0, arrow_xy$x, arrow_xy$y,
                       length = 0.08,
                       lwd = 2.0,
                       col = MODE_COLORS[mode])

                points(arrow_xy$x, arrow_xy$y,
                       pch = 16,
                       cex = 0.7,
                       col = MODE_COLORS[mode])

                arrow_endpoints[[mode]] = c(x = arrow_xy$x, y = arrow_xy$y)
            }
        }
    }

    if (star != "" &&
        !is.null(arrow_endpoints$swim) &&
        !is.null(arrow_endpoints$walk)) {

        star_x = mean(c(arrow_endpoints$swim["x"], arrow_endpoints$walk["x"]))
        star_y = mean(c(arrow_endpoints$swim["y"], arrow_endpoints$walk["y"]))

        star_distance = sqrt(star_x^2 + star_y^2)

        if (is.finite(star_distance) && star_distance > 0) {
            star_x = star_x + SIGNIFICANCE_OFFSET * star_x / star_distance
            star_y = star_y + SIGNIFICANCE_OFFSET * star_y / star_distance
        }

        text(star_x, star_y,
             labels = star,
             cex = SIGNIFICANCE_CEX,
             font = 2)
    }
}

plot_phase_circular = function(relative_phase, phase_comparisons) {
    output_path = file.path(PLOT_DIR, "swim_walk_relative_phase_circular.pdf")

    pair_order = combn(LEGS, 2, simplify = FALSE)
    pair_order = vapply(pair_order,
                        function(pair) paste0(pair[1], "-", pair[2]),
                        character(1))

    panel_labels = LETTERS[seq_along(pair_order)]

    pdf(output_path, width = 14, height = 8.5)

    par(mfrow = c(3, 5),
        mar = c(1.4, 1.4, 1.8, 1.1),
        oma = c(0.5, 0.5, 1.8, 0.5))

    for (i in seq_along(pair_order)) {
        pair_name = pair_order[i]
        comparison_row = phase_comparisons[phase_comparisons$pair == pair_name, ]

        draw_phase_circle(pair_name = pair_name,
                          relative_phase = relative_phase,
                          comparison_row = comparison_row,
                          panel_label = panel_labels[i])
    }

    dev.off()
    return(output_path)
}


relative_phase = read_data(file.path(RELATIVE_PHASE_DIR, "relative_phase.csv"))
phase_comparisons = read_data(file.path(PHASE_STAT_DIR, "swim_walk_phase_comparisons.csv"))

plot_phase_circular(relative_phase, phase_comparisons)
```
