## Supplemental Methods for "Data-driven frame-by-frame analysis of surface swimming and terrestrial walking in the coastal beetle *Phaleria atriceps*": 07_six-leg_phase_configuration_analysis.R

```
PHASE_DIR = file.path("scratch_results", "phase_analysis", "recordings")
OUTPUT_DIR = file.path("scratch_results", "six_leg_phase_configuration")
DATA_DIR = file.path(OUTPUT_DIR, "recordings")
PLOT_DIR = file.path(OUTPUT_DIR, "plot")

LEGS = c("L1", "L2", "L3", "R1", "R2", "R3")
MODES = c("swim", "walk")
BOOTSTRAP_N = 5000
BOOTSTRAP_SEED = 20261006
MAX_BOOTSTRAP_ATTEMPT_FACTOR = 5
PAIR_WEIGHT_METHOD = "equal"
WRITE_BOOTSTRAP_REPLICATES = TRUE
WRITE_BOOTSTRAP_ATTEMPT_LOG = TRUE

dir.create(OUTPUT_DIR, recursive = TRUE, showWarnings = FALSE)
dir.create(DATA_DIR, recursive = TRUE, showWarnings = FALSE)
dir.create(PLOT_DIR, recursive = TRUE, showWarnings = FALSE)

pair_list = combn(LEGS, 2, simplify = FALSE)
PAIR_TABLE = do.call(rbind, lapply(pair_list, function(x) {
    data.frame(pair = paste0(x[1], "-", x[2]), leg1 = x[1], leg2 = x[2], stringsAsFactors = FALSE)
}))
rownames(PAIR_TABLE) = NULL
read_data = function(path) {
    read.csv(path, header = TRUE, stringsAsFactors = FALSE, check.names = FALSE, na.strings = c("", "NA", "NaN", "nan"))
}
wrap_pi = function(angle) atan2(sin(angle), cos(angle))
rad_to_deg = function(angle) 180 * angle / pi
rad_to_deg_360 = function(angle) {
    degree = (180 * angle / pi) %% 360
    degree[degree < 0] = degree[degree < 0] + 360
    degree
}
circular_mean = function(angle) {
    angle = angle[is.finite(angle)]
    if (length(angle) == 0) return(NA_real_)
    atan2(mean(sin(angle)), mean(cos(angle)))
}
resultant_length = function(angle) {
    angle = angle[is.finite(angle)]
    if (length(angle) == 0) return(NA_real_)
    sqrt(mean(sin(angle))^2 + mean(cos(angle))^2)
}
unwrap_to_reference = function(angle, reference) reference + wrap_pi(angle - reference)
validate_relative_phase = function(data) {
    required_columns = c("id", "mode", "pair", "leg1", "leg2", "relative_phase_rad")
    missing_columns = setdiff(required_columns, names(data))
    if (length(missing_columns) > 0) stop("Missing columns: ", paste(missing_columns, collapse = ", "))
    if (any(!data$mode %in% MODES)) stop("Unexpected mode values.")
    if (any(!data$pair %in% PAIR_TABLE$pair)) stop("Unexpected leg pair names.")
    if (any(duplicated(data[, c("id", "pair")]))) stop("Duplicated id x pair observations were detected.")
    invisible(TRUE)
}
make_pair_means = function(data, mode_name) {
    result = vector("list", nrow(PAIR_TABLE))

    for (k in seq_len(nrow(PAIR_TABLE))) {
        pair_name = PAIR_TABLE$pair[k]
        z = data[data$mode == mode_name & data$pair == pair_name & is.finite(data$relative_phase_rad), , drop = FALSE]
        if (nrow(z) == 0) next

        mean_phase = circular_mean(z$relative_phase_rad)
        result[[k]] = data.frame(
            mode = mode_name, pair = pair_name, leg1 = PAIR_TABLE$leg1[k], leg2 = PAIR_TABLE$leg2[k],
            n = nrow(z), mean_phase_rad = mean_phase, mean_phase_deg_signed = rad_to_deg(mean_phase),
            mean_phase_deg_360 = rad_to_deg_360(mean_phase),
            resultant_length_R = resultant_length(z$relative_phase_rad),
            stringsAsFactors = FALSE
        )
    }

    keep = !vapply(result, is.null, logical(1))
    if (!any(keep)) return(data.frame())

    result = do.call(rbind, result[keep])
    rownames(result) = NULL
    result
}

make_pair_weights = function(pair_mean) {
    if (PAIR_WEIGHT_METHOD == "equal") {
        weight = rep(1, nrow(pair_mean))
    } else if (PAIR_WEIGHT_METHOD == "n") {
        weight = pair_mean$n
    } else if (PAIR_WEIGHT_METHOD == "nR") {
        weight = pair_mean$n * pair_mean$resultant_length_R
    } else {
        stop("Unknown PAIR_WEIGHT_METHOD: ", PAIR_WEIGHT_METHOD)
    }

    if (any(!is.finite(weight)) || all(weight <= 0)) stop("Invalid pair weights.")
    weight[weight < 0] = 0
    weight = weight / mean(weight[weight > 0])
    weight
}
configuration_is_connected = function(pair_mean) {
    adjacency = matrix(FALSE, nrow = length(LEGS), ncol = length(LEGS), dimnames = list(LEGS, LEGS))
    for (k in seq_len(nrow(pair_mean))) {
        leg1 = pair_mean$leg1[k]
        leg2 = pair_mean$leg2[k]
        adjacency[leg1, leg2] = TRUE
        adjacency[leg2, leg1] = TRUE
    }
    visited = setNames(rep(FALSE, length(LEGS)), LEGS)
    queue = LEGS[1]
    visited[LEGS[1]] = TRUE
    while (length(queue) > 0) {
        current = queue[1]
        queue = if (length(queue) == 1) character(0) else queue[-1]
        neighbours = LEGS[adjacency[current, ]]
        new_neighbours = neighbours[!visited[neighbours]]

        if (length(new_neighbours) > 0) {
            visited[new_neighbours] = TRUE
            queue = c(queue, new_neighbours)
        }
    }
    all(visited)
}
configuration_status = function(pair_mean) {
    n_pairs = nrow(pair_mean)

    if (n_pairs < length(LEGS) - 1) {
        return(list(n_pairs = n_pairs, connected = FALSE, fit_possible = FALSE, reason = "too_few_pairs"))
    }

    connected = configuration_is_connected(pair_mean)

    if (!connected) {
        return(list(n_pairs = n_pairs, connected = FALSE, fit_possible = FALSE, reason = "disconnected"))
    }

    list(n_pairs = n_pairs, connected = TRUE, fit_possible = TRUE, reason = "ok")
}

get_pair_presence = function(data) {
    observed_pairs = unique(data$pair[is.finite(data$relative_phase_rad)])
    presence = PAIR_TABLE$pair %in% observed_pairs
    names(presence) = PAIR_TABLE$pair
    presence
}

make_initial_configuration = function(pair_mean) {
    phase = setNames(rep(NA_real_, length(LEGS)), LEGS)
    phase["L1"] = 0

    repeat {
        changed = FALSE

        for (k in seq_len(nrow(pair_mean))) {
            leg1 = pair_mean$leg1[k]
            leg2 = pair_mean$leg2[k]
            difference = pair_mean$mean_phase_rad[k]

            if (is.finite(phase[leg1]) && !is.finite(phase[leg2])) {
                phase[leg2] = phase[leg1] + difference
                changed = TRUE
            }

            if (is.finite(phase[leg2]) && !is.finite(phase[leg1])) {
                phase[leg1] = phase[leg2] - difference
                changed = TRUE
            }
        }

        if (!changed) break
    }

    if (any(!is.finite(phase))) return(NULL)

    phase = wrap_pi(phase - phase["L1"])
    phase["L1"] = 0
    phase
}
fit_phase_configuration = function(pair_mean, start_configuration = NULL) {
    if (nrow(pair_mean) < length(LEGS) - 1) return(NULL)
    if (!configuration_is_connected(pair_mean)) return(NULL)

    initial = make_initial_configuration(pair_mean)
    if (is.null(initial)) return(NULL)

    design = matrix(0, nrow = nrow(pair_mean), ncol = length(LEGS), dimnames = list(NULL, LEGS))

    for (k in seq_len(nrow(pair_mean))) {
        design[k, pair_mean$leg1[k]] = -1
        design[k, pair_mean$leg2[k]] = 1
    }

    observed = pair_mean$mean_phase_rad
    weight = make_pair_weights(pair_mean)

    objective = function(parameter) {
        configuration = c(0, parameter)
        names(configuration) = LEGS
        predicted = drop(design %*% configuration)
        residual = wrap_pi(observed - predicted)
        sum(weight * (1 - cos(residual)))
    }

    starts = list(initial[-1], rep(0, length(LEGS) - 1))

    if (!is.null(start_configuration)) {
        start_configuration = start_configuration[LEGS]
        start_configuration = wrap_pi(start_configuration - start_configuration["L1"])
        starts = c(list(start_configuration[-1]), starts)
    }

    candidate_fits = vector("list", length(starts))

    for (k in seq_along(starts)) {
        candidate_fits[[k]] = tryCatch(
            optim(par = starts[[k]], fn = objective, method = "BFGS",
                  control = list(maxit = 2000, reltol = 1e-12)),
            error = function(e) NULL
        )
    }

    valid_fit = vapply(candidate_fits, function(x) !is.null(x) && is.finite(x$value), logical(1))
    candidate_fits = candidate_fits[valid_fit]

    if (length(candidate_fits) == 0) {
        fallback = tryCatch(
            optim(par = initial[-1], fn = objective, method = "Nelder-Mead",
                  control = list(maxit = 5000, reltol = 1e-12)),
            error = function(e) NULL
        )

        if (is.null(fallback) || !is.finite(fallback$value)) return(NULL)
        best_fit = fallback
    } else {
        objective_values = vapply(candidate_fits, function(x) x$value, numeric(1))
        best_fit = candidate_fits[[which.min(objective_values)]]
    }

    configuration = c(0, best_fit$par)
    names(configuration) = LEGS
    configuration = wrap_pi(configuration - configuration["L1"])
    configuration["L1"] = 0

    predicted = wrap_pi(drop(design %*% configuration))
    residual = wrap_pi(observed - predicted)
    residual_deg = rad_to_deg(residual)

    list(
        phase_rad = configuration,
        predicted_pair_phase_rad = predicted,
        residual_rad = residual,
        residual_rmse_deg = sqrt(mean(residual_deg^2)),
        residual_mae_deg = mean(abs(residual_deg)),
        objective = best_fit$value,
        convergence = best_fit$convergence
    )
}
fit_mode = function(data, mode_name, start_configuration = NULL) {
    pair_mean = make_pair_means(data, mode_name)
    fit = fit_phase_configuration(pair_mean, start_configuration)
    if (is.null(fit)) return(NULL)
    list(pair_mean = pair_mean, fit = fit)
}
align_mode_configurations = function(walk_phase, swim_phase) {
    walk_phase = walk_phase[LEGS]
    swim_phase = swim_phase[LEGS]

    raw_difference = wrap_pi(swim_phase - walk_phase)
    common_rotation = circular_mean(raw_difference)
    alignment_R = resultant_length(raw_difference)

    swim_aligned = wrap_pi(swim_phase - common_rotation)
    phase_shift = wrap_pi(swim_aligned - walk_phase)

    names(swim_aligned) = LEGS
    names(phase_shift) = LEGS

    list(
        common_rotation_rad = common_rotation,
        alignment_R = alignment_R,
        walk_phase_rad = walk_phase,
        swim_phase_aligned_rad = swim_aligned,
        phase_shift_rad = phase_shift
    )
}

make_reconstruction_table = function(mode_fit, mode_name) {
    result = mode_fit$pair_mean
    result$fitted_phase_rad = mode_fit$fit$predicted_pair_phase_rad
    result$fitted_phase_deg_signed = rad_to_deg(result$fitted_phase_rad)
    result$fitted_phase_deg_360 = rad_to_deg_360(result$fitted_phase_rad)
    result$residual_rad = mode_fit$fit$residual_rad
    result$residual_deg = rad_to_deg(result$residual_rad)
    result$mode = mode_name
    result
}

resample_mode_by_individual = function(data, mode_name) {
    z = data[data$mode == mode_name, , drop = FALSE]
    individuals = unique(z$id)
    sampled = sample(individuals, length(individuals), replace = TRUE)

    result = lapply(seq_along(sampled), function(k) {
        d = z[z$id == sampled[k], , drop = FALSE]
        d$id = paste0(mode_name, "_bootstrap_", k)
        d
    })

    result = do.call(rbind, result)
    rownames(result) = NULL
    result
}

relative_phase = read_data(file.path(PHASE_DIR, "relative_phase.csv"))
validate_relative_phase(relative_phase)
relative_phase = relative_phase[is.finite(relative_phase$relative_phase_rad), , drop = FALSE]
walk_result = fit_mode(relative_phase, "walk")
swim_result = fit_mode(relative_phase, "swim")
if (is.null(walk_result) || is.null(swim_result)) {
    stop("The six leg phase configuration could not be reconstructed.")
}
walk_phase = walk_result$fit$phase_rad
swim_phase = swim_result$fit$phase_rad
alignment = align_mode_configurations(walk_phase, swim_phase)
pair_means = rbind(walk_result$pair_mean, swim_result$pair_mean)
write.csv(pair_means, file.path(DATA_DIR, "mode_pair_mean_phase.csv"), row.names = FALSE)
configuration_L1 = rbind(
    data.frame(mode = "walk", leg = LEGS, phase_rad = walk_phase[LEGS],
               phase_deg_signed = rad_to_deg(walk_phase[LEGS]),
               phase_deg_360 = rad_to_deg_360(walk_phase[LEGS]), stringsAsFactors = FALSE),
    data.frame(mode = "swim", leg = LEGS, phase_rad = swim_phase[LEGS],
               phase_deg_signed = rad_to_deg(swim_phase[LEGS]),
               phase_deg_360 = rad_to_deg_360(swim_phase[LEGS]), stringsAsFactors = FALSE)
)
write.csv(configuration_L1, file.path(DATA_DIR, "mode_leg_phase_configuration_L1_reference.csv"), row.names = FALSE)

configuration_aligned = rbind(
    data.frame(mode = "walk", leg = LEGS, phase_rad = alignment$walk_phase_rad[LEGS],
               phase_deg_signed = rad_to_deg(alignment$walk_phase_rad[LEGS]),
               phase_deg_360 = rad_to_deg_360(alignment$walk_phase_rad[LEGS]), stringsAsFactors = FALSE),
    data.frame(mode = "swim", leg = LEGS, phase_rad = alignment$swim_phase_aligned_rad[LEGS],
               phase_deg_signed = rad_to_deg(alignment$swim_phase_aligned_rad[LEGS]),
               phase_deg_360 = rad_to_deg_360(alignment$swim_phase_aligned_rad[LEGS]), stringsAsFactors = FALSE)
)
write.csv(configuration_aligned, file.path(DATA_DIR, "mode_leg_phase_configuration_walk_aligned.csv"), row.names = FALSE)

walk_reconstruction = make_reconstruction_table(walk_result, "walk")
swim_reconstruction = make_reconstruction_table(swim_result, "swim")
reconstruction = rbind(walk_reconstruction, swim_reconstruction)

write.csv(reconstruction, file.path(DATA_DIR, "pair_phase_reconstruction.csv"), row.names = FALSE)

set.seed(BOOTSTRAP_SEED)

bootstrap_shift = matrix(NA_real_, nrow = BOOTSTRAP_N, ncol = length(LEGS), dimnames = list(NULL, LEGS))

missing_attempt_walk = setNames(integer(nrow(PAIR_TABLE)), PAIR_TABLE$pair)
missing_attempt_swim = setNames(integer(nrow(PAIR_TABLE)), PAIR_TABLE$pair)
missing_success_walk = setNames(integer(nrow(PAIR_TABLE)), PAIR_TABLE$pair)
missing_success_swim = setNames(integer(nrow(PAIR_TABLE)), PAIR_TABLE$pair)

failure_counts = c(
    walk_too_few_pairs = 0L,
    walk_disconnected = 0L,
    walk_optimization_failure = 0L,
    swim_too_few_pairs = 0L,
    swim_disconnected = 0L,
    swim_optimization_failure = 0L
)

success = 0
attempt = 0
MAX_ATTEMPT = BOOTSTRAP_N * MAX_BOOTSTRAP_ATTEMPT_FACTOR
attempt_log = vector("list", MAX_ATTEMPT)

while (success < BOOTSTRAP_N && attempt < MAX_ATTEMPT) {
    attempt = attempt + 1

    walk_boot = resample_mode_by_individual(relative_phase, "walk")
    swim_boot = resample_mode_by_individual(relative_phase, "swim")

    walk_presence = get_pair_presence(walk_boot)
    swim_presence = get_pair_presence(swim_boot)

    missing_attempt_walk = missing_attempt_walk + as.integer(!walk_presence)
    missing_attempt_swim = missing_attempt_swim + as.integer(!swim_presence)

    walk_pair_mean = make_pair_means(walk_boot, "walk")
    swim_pair_mean = make_pair_means(swim_boot, "swim")

    walk_status = configuration_status(walk_pair_mean)
    swim_status = configuration_status(swim_pair_mean)

    walk_fit_object = NULL
    swim_fit_object = NULL

    if (!walk_status$fit_possible) {
        if (walk_status$reason == "too_few_pairs") failure_counts["walk_too_few_pairs"] = failure_counts["walk_too_few_pairs"] + 1L
        if (walk_status$reason == "disconnected") failure_counts["walk_disconnected"] = failure_counts["walk_disconnected"] + 1L
    } else {
        walk_fit_object = fit_phase_configuration(walk_pair_mean, start_configuration = walk_phase)
        if (is.null(walk_fit_object)) failure_counts["walk_optimization_failure"] = failure_counts["walk_optimization_failure"] + 1L
    }

    if (!swim_status$fit_possible) {
        if (swim_status$reason == "too_few_pairs") failure_counts["swim_too_few_pairs"] = failure_counts["swim_too_few_pairs"] + 1L
        if (swim_status$reason == "disconnected") failure_counts["swim_disconnected"] = failure_counts["swim_disconnected"] + 1L
    } else {
        swim_fit_object = fit_phase_configuration(swim_pair_mean, start_configuration = swim_phase)
        if (is.null(swim_fit_object)) failure_counts["swim_optimization_failure"] = failure_counts["swim_optimization_failure"] + 1L
    }

    walk_fit_success = !is.null(walk_fit_object)
    swim_fit_success = !is.null(swim_fit_object)
    replicate_success = walk_fit_success && swim_fit_success

    attempt_log[[attempt]] = data.frame(
        attempt = attempt,
        n_walk_pairs = walk_status$n_pairs,
        n_swim_pairs = swim_status$n_pairs,
        walk_connected = walk_status$connected,
        swim_connected = swim_status$connected,
        walk_fit_success = walk_fit_success,
        swim_fit_success = swim_fit_success,
        replicate_success = replicate_success,
        stringsAsFactors = FALSE
    )

    if (!replicate_success) next

    success = success + 1

    missing_success_walk = missing_success_walk + as.integer(!walk_presence)
    missing_success_swim = missing_success_swim + as.integer(!swim_presence)

    boot_alignment = align_mode_configurations(
        walk_phase = walk_fit_object$phase_rad,
        swim_phase = swim_fit_object$phase_rad
    )

    bootstrap_shift[success, ] = unwrap_to_reference(
        boot_alignment$phase_shift_rad[LEGS],
        alignment$phase_shift_rad[LEGS]
    )
}

if (success == 0) stop("No bootstrap replicate was successful.")

if (success < BOOTSTRAP_N) {
    warning("Only ", success, " of ", BOOTSTRAP_N, " requested bootstrap replicates were successful.")
    bootstrap_shift = bootstrap_shift[seq_len(success), , drop = FALSE]
}

attempt_log = do.call(rbind, attempt_log[seq_len(attempt)])
failed_attempts = attempt - success
failure_rate = failed_attempts / attempt

point_shift_rad = alignment$phase_shift_rad[LEGS]
point_shift_deg = rad_to_deg(point_shift_rad)
bootstrap_shift_deg = rad_to_deg(bootstrap_shift)

lower_95 = apply(bootstrap_shift_deg, 2, quantile, probs = 0.025, na.rm = TRUE, type = 7)
upper_95 = apply(bootstrap_shift_deg, 2, quantile, probs = 0.975, na.rm = TRUE, type = 7)

point_matrix = matrix(point_shift_deg, nrow = nrow(bootstrap_shift_deg), ncol = length(LEGS), byrow = TRUE)
bootstrap_error_deg = bootstrap_shift_deg - point_matrix
max_absolute_error = apply(abs(bootstrap_error_deg), 1, max, na.rm = TRUE)

simultaneous_margin_95 = unname(
    quantile(max_absolute_error, probs = 0.95, na.rm = TRUE, type = 7)
)

shift_summary = data.frame(
    leg = LEGS,
    phase_shift_rad = point_shift_rad,
    phase_shift_deg = point_shift_deg,
    magnitude_deg = abs(point_shift_deg),
    cycle_fraction = point_shift_deg / 360,
    magnitude_percent_cycle = 100 * abs(point_shift_deg) / 360,
    lower_95_deg = lower_95[LEGS],
    upper_95_deg = upper_95[LEGS],
    simultaneous_lower_95_deg = point_shift_deg - simultaneous_margin_95,
    simultaneous_upper_95_deg = point_shift_deg + simultaneous_margin_95,
    direction = ifelse(point_shift_deg > 0, "phase advance",
                       ifelse(point_shift_deg < 0, "phase delay", "no change")),
    pointwise_CI_excludes_zero = lower_95[LEGS] > 0 | upper_95[LEGS] < 0,
    simultaneous_CI_excludes_zero =
        (point_shift_deg - simultaneous_margin_95) > 0 |
        (point_shift_deg + simultaneous_margin_95) < 0,
    stringsAsFactors = FALSE
)

write.csv(shift_summary, file.path(DATA_DIR, "walk_to_swim_leg_phase_shift.csv"), row.names = FALSE)
if (WRITE_BOOTSTRAP_REPLICATES) {
    bootstrap_output = data.frame(
        bootstrap = seq_len(nrow(bootstrap_shift_deg)),
        bootstrap_shift_deg,
        check.names = FALSE
    )

    write.csv(bootstrap_output, file.path(DATA_DIR, "walk_to_swim_leg_phase_shift_bootstrap.csv"), row.names = FALSE)
}

if (WRITE_BOOTSTRAP_ATTEMPT_LOG) {
    write.csv(attempt_log, file.path(DATA_DIR, "bootstrap_attempt_log.csv"), row.names = FALSE)
}
original_walk_n = setNames(rep(0L, nrow(PAIR_TABLE)), PAIR_TABLE$pair)
original_swim_n = setNames(rep(0L, nrow(PAIR_TABLE)), PAIR_TABLE$pair)

original_walk_n[walk_result$pair_mean$pair] = walk_result$pair_mean$n
original_swim_n[swim_result$pair_mean$pair] = swim_result$pair_mean$n

pair_missingness = rbind(
    data.frame(
        mode = "walk",
        pair = PAIR_TABLE$pair,
        original_n = original_walk_n[PAIR_TABLE$pair],
        missing_attempts = missing_attempt_walk[PAIR_TABLE$pair],
        missing_rate_all_attempts = missing_attempt_walk[PAIR_TABLE$pair] / attempt,
        missing_successful_replicates = missing_success_walk[PAIR_TABLE$pair],
        missing_rate_successful_replicates = missing_success_walk[PAIR_TABLE$pair] / success,
        stringsAsFactors = FALSE
    ),
    data.frame(
        mode = "swim",
        pair = PAIR_TABLE$pair,
        original_n = original_swim_n[PAIR_TABLE$pair],
        missing_attempts = missing_attempt_swim[PAIR_TABLE$pair],
        missing_rate_all_attempts = missing_attempt_swim[PAIR_TABLE$pair] / attempt,
        missing_successful_replicates = missing_success_swim[PAIR_TABLE$pair],
        missing_rate_successful_replicates = missing_success_swim[PAIR_TABLE$pair] / success,
        stringsAsFactors = FALSE
    )
)

rownames(pair_missingness) = NULL
write.csv(pair_missingness, file.path(DATA_DIR, "bootstrap_pair_missingness.csv"), row.names = FALSE)

failure_reason_table = data.frame(
    reason = names(failure_counts),
    count = as.integer(failure_counts),
    rate_per_attempt = as.integer(failure_counts) / attempt,
    stringsAsFactors = FALSE
)

write.csv(failure_reason_table, file.path(DATA_DIR, "bootstrap_failure_reasons.csv"), row.names = FALSE)

diagnostics = data.frame(
    quantity = c(
        "walk_pair_reconstruction_RMSE_deg",
        "walk_pair_reconstruction_MAE_deg",
        "swim_pair_reconstruction_RMSE_deg",
        "swim_pair_reconstruction_MAE_deg",
        "walk_circular_objective",
        "swim_circular_objective",
        "mode_alignment_resultant_length_R",
        "removed_common_rotation_deg",
        "bootstrap_requested_successful_replicates",
        "bootstrap_successful_replicates",
        "bootstrap_total_attempts",
        "bootstrap_failed_attempts",
        "bootstrap_failure_rate",
        "simultaneous_95_margin_deg"
    ),
    value = c(
        walk_result$fit$residual_rmse_deg,
        walk_result$fit$residual_mae_deg,
        swim_result$fit$residual_rmse_deg,
        swim_result$fit$residual_mae_deg,
        walk_result$fit$objective,
        swim_result$fit$objective,
        alignment$alignment_R,
        rad_to_deg(alignment$common_rotation_rad),
        BOOTSTRAP_N,
        success,
        attempt,
        failed_attempts,
        failure_rate,
        simultaneous_margin_95
    ),
    stringsAsFactors = FALSE
)

write.csv(diagnostics, file.path(DATA_DIR, "phase_configuration_diagnostics.csv"), row.names = FALSE)

plot_configuration = function(phase, main) {
    plot(NA, NA, xlim = c(-1.35, 1.35), ylim = c(-1.35, 1.35), asp = 1,
         axes = FALSE, xlab = "", ylab = "", bty = "n", main = main)

    theta = seq(0, 2*pi, length.out = 361)
    lines(cos(theta), sin(theta))
    segments(-1, 0, 1, 0, col = "grey75")
    segments(0, -1, 0, 1, col = "grey75")

    text(1.15, 0, "0°")
    text(0, 1.15, "90°")
    text(-1.15, 0, "180°")
    text(0, -1.15, "270°")

    for (leg in LEGS) {
        angle = phase[leg]
        x = cos(angle)
        y = sin(angle)

        segments(0, 0, x, y, lwd = 1.5)
        points(x, y, pch = 16, cex = 1.1)
        text(1.12*x, 1.12*y, labels = leg, font = 2)
    }
}

pdf(file.path(PLOT_DIR, "mode_six_leg_phase_configuration.pdf"), width = 9, height = 4.5)
par(mfrow = c(1, 2), mar = c(1, 1, 2.5, 1))
plot_configuration(walk_phase, "Walking")
plot_configuration(swim_phase, "Swimming")
dev.off()

pdf(file.path(PLOT_DIR, "walk_to_swim_leg_phase_shift.pdf"), width = 7, height = 5.5)
par(mar = c(5, 5, 1.5, 1), las = 1, tck = 0.015)

x = seq_along(LEGS)
ylim = range(c(shift_summary$simultaneous_lower_95_deg,
               shift_summary$simultaneous_upper_95_deg, 0), finite = TRUE)

plot(x, shift_summary$phase_shift_deg, type = "n", xaxt = "n",
     xlab = "Leg", ylab = "Relative phase shift from walking to swimming (deg)",
     ylim = ylim, bty = "n")

axis(1, at = x, labels = LEGS)
abline(h = 0, lty = 2)

segments(x, shift_summary$simultaneous_lower_95_deg,
         x, shift_summary$simultaneous_upper_95_deg, lwd = 1)

segments(x, shift_summary$lower_95_deg,
         x, shift_summary$upper_95_deg, lwd = 3)

points(x, shift_summary$phase_shift_deg, pch = 16, cex = 1.2)
dev.off()

metadata = c(
    "Six leg phase configuration analysis",
    "",
    "Input:",
    "- relative_phase.csv",
    "- one representative relative phase per individual and anatomical leg pair",
    "- each pair retains its original pair specific overlap midpoint",
    "",
    "Mode summary:",
    "- circular mean calculated separately for each anatomical leg pair",
    "",
    "Six leg reconstruction:",
    "- observed pair phase = phase of leg2 - phase of leg1",
    "- six latent leg phases fitted simultaneously",
    "- circular loss = sum w * [1 - cos(observed - fitted)]",
    paste0("- pair weighting method = ", PAIR_WEIGHT_METHOD),
    "- L1 fixed to 0 degrees only to define the angular origin within each mode",
    "",
    "Walking and swimming comparison:",
    "- arbitrary common rotation between configurations removed before calculating leg specific changes",
    "- positive value = relative phase advance during swimming",
    "- negative value = relative phase delay during swimming",
    "",
    "Bootstrap:",
    "- resampling unit = individual",
    "- individuals resampled independently within walking and swimming",
    "- all available pairwise phase values belonging to a sampled individual retained together",
    "- missing pair information within an individual retained as observed",
    "- pairwise circular means recalculated in every replicate",
    "- six leg configuration reconstructed in every replicate",
    "- configuration alignment repeated in every replicate",
    "- requested successful bootstrap replicates = 5000",
    "- unsuccessful attempts excluded from the bootstrap distribution",
    "",
    "Bootstrap diagnostics:",
    "- bootstrap_attempt_log.csv records pair availability, connectivity, and fit success for each attempt",
    "- bootstrap_pair_missingness.csv records how often each anatomical pair was absent after individual resampling",
    "- bootstrap_failure_reasons.csv records failures caused by too few pairs, disconnected leg networks, or fitting failure",
    "- phase_configuration_diagnostics.csv records total attempts, successful replicates, failed attempts, and failure rate"
)

writeLines(metadata, file.path(OUTPUT_DIR, "analysis_metadata.txt"))

cat("\n============================================================\n")
cat("Mode specific six leg phase configuration\n")
cat("L1 = 0 degrees within each mode\n")
cat("============================================================\n\n")
print(configuration_L1)

cat("\n============================================================\n")
cat("Walking to swimming relative phase changes\n")
cat("Positive = relative phase advance\n")
cat("Negative = relative phase delay\n")
cat("============================================================\n\n")
print(shift_summary)

cat("\n============================================================\n")
cat("Reconstruction and bootstrap diagnostics\n")
cat("============================================================\n\n")
print(diagnostics)

cat("\n============================================================\n")
cat("Bootstrap failure reasons\n")
cat("============================================================\n\n")
print(failure_reason_table)

cat("\n============================================================\n")
cat("Pair missingness during bootstrap\n")
cat("============================================================\n\n")
print(pair_missingness)

cat("\nOutput directory:\n")
cat(OUTPUT_DIR, "\n")
```
