## Supplemental Methods for "Data-driven frame-by-frame analysis of surface swimming and terrestrial walking in the coastal beetle *Phaleria atriceps*": 07b_plot_six-leg_phase_configuration.R

```
INPUT_DIR = file.path("scratch_results", "six_leg_phase_configuration", "recordings")
OUTPUT_DIR = file.path("scratch_results", "six_leg_phase_configuration", "plot")

CONFIG_L1_FILE = file.path(INPUT_DIR, "mode_leg_phase_configuration_L1_reference.csv")
CONFIG_ALIGNED_FILE = file.path(INPUT_DIR, "mode_leg_phase_configuration_walk_aligned.csv")
SHIFT_FILE = file.path(INPUT_DIR, "walk_to_swim_leg_phase_shift.csv")

OUTPUT_PDF = file.path(OUTPUT_DIR, "six_leg_phase_configuration_AB.pdf")
OUTPUT_PNG = file.path(OUTPUT_DIR, "six_leg_phase_configuration_AB.png")

LEGS = c("L1", "L2", "L3", "R1", "R2", "R3")

# Plot settings
FONT_FAMILY = "Arial"
POINT_CEX = 1.15
ERROR_CAP_WIDTH = 0.10
MODE_OFFSET = 0.11

# Walking and swimming are distinguished by symbol/fill rather than color.
WALK_PCH = 16
SWIM_PCH = 16
WALK_COLOR = "#D62728"
SWIM_COLOR = "#1F77B4"

dir.create(OUTPUT_DIR, recursive = TRUE, showWarnings = FALSE)

read_data = function(path) {
    if (!file.exists(path)) stop("Input file not found: ", path)
    read.csv(path, header = TRUE, stringsAsFactors = FALSE, check.names = FALSE,
             na.strings = c("", "NA", "NaN", "nan"))
}

validate_columns = function(data, required, file_name) {
    missing_columns = setdiff(required, names(data))
    if (length(missing_columns) > 0) {
        stop("Missing columns in ", file_name, ": ", paste(missing_columns, collapse = ", "))
    }
}

phase_to_plot_range = function(x) {
    y = x %% 360
    y[y < 0] = y[y < 0] + 360
    y[y > 270] = y[y > 270] - 360
    y
}

# Draw a vertical confidence interval with short horizontal caps.
draw_ci = function(x, lower, upper, cap_width = ERROR_CAP_WIDTH, lwd = 1.5) {
    segments(x0 = x, y0 = lower, x1 = x, y1 = upper, lwd = lwd)
    segments(x0 = x - cap_width, y0 = lower, x1 = x + cap_width, y1 = lower, lwd = lwd)
    segments(x0 = x - cap_width, y0 = upper, x1 = x + cap_width, y1 = upper, lwd = lwd)
}

plot_panel_single_mode = function(configuration, mode_name, panel_label) {
    z = configuration[configuration$mode == mode_name, , drop = FALSE]
    z = z[match(LEGS, z$leg), , drop = FALSE]

    x = seq_along(LEGS)

    plot(NA, NA,
         xlim = c(0.5, length(LEGS) + 0.5),
         ylim = c(-90, 270),
         xaxt = "n", yaxt = "n",
         xlab = "Leg",
         ylab = "Six-leg phase relationship (deg)",
         bty = "n")

    axis(1, at = x, labels = LEGS)
    axis(2, at = seq(-90, 270, by = 90), labels = seq(-90, 270, by = 90), las = 1)

    abline(h = c(0, 180), lty = 2, lwd = 1)

    if (mode_name == "walk") {
        points(x, z$phase_deg_plot,
               pch = WALK_PCH,
               col = WALK_COLOR, cex = POINT_CEX)
    } else if (mode_name == "swim") {
        points(x, z$phase_deg_plot,
               pch = SWIM_PCH, col = SWIM_COLOR,
               cex = POINT_CEX)
    } else {
        stop("Unknown mode_name: ", mode_name)
    }

    mtext(panel_label, side = 3, adj = 0, line = 0.35, font = 2, cex = 2.0)
}

plot_panel_aligned = function(configuration) {
    walk = configuration[configuration$mode == "walk", , drop = FALSE]
    swim = configuration[configuration$mode == "swim", , drop = FALSE]

    walk = walk[match(LEGS, walk$leg), , drop = FALSE]
    swim = swim[match(LEGS, swim$leg), , drop = FALSE]

    x = seq_along(LEGS)
    x_walk = x - MODE_OFFSET
    x_swim = x + MODE_OFFSET

    plot(NA, NA,
         xlim = c(0.5, length(LEGS) + 0.5),
         ylim = c(-90, 270),
         xaxt = "n", yaxt = "n",
         xlab = "Leg",
         ylab = "Six-leg phase relationship (deg)",
         bty = "n")

    axis(1, at = x, labels = LEGS)
    axis(2, at = seq(-90, 270, by = 90), labels = seq(-90, 270, by = 90), las = 1)

    abline(h = c(0, 180), lty = 2, lwd = 1)

    points(x_walk, walk$phase_deg_plot,
           pch = WALK_PCH, 
           col = WALK_COLOR, cex = POINT_CEX)
    points(x_swim, swim$phase_deg_plot,
           pch = SWIM_PCH, col = SWIM_COLOR,
           cex = POINT_CEX)

    mtext("C", side = 3, adj = 0, line = 0.35, font = 2, cex = 2.0)
}

plot_panel_shift = function(shift) {
    shift = shift[match(LEGS, shift$leg), , drop = FALSE]
    x = seq_along(LEGS)

    y_min = min(c(shift$lower_95_deg, 0), na.rm = TRUE)
    y_max = max(c(shift$upper_95_deg, 0), na.rm = TRUE)

    # Round the limits outward to 30-degree increments for a clean axis.
    y_min = floor(y_min / 30) * 30
    y_max = ceiling(y_max / 30) * 30

    if (y_min == y_max) {
        y_min = y_min - 30
        y_max = y_max + 30
    }

    plot(NA, NA,
         xlim = c(0.5, length(LEGS) + 0.5),
         ylim = c(y_min, y_max),
         xaxt = "n", yaxt = "n",
         xlab = "Leg",
         ylab = "Phase differences\nbetween walking and swimming (deg)",
         bty = "n")

    axis(1, at = x, labels = LEGS)
    y_ticks = seq(y_min, y_max, by = 30)
    axis(2, at = y_ticks, labels = y_ticks, las = 1)

    abline(h = 0, lty = 2, lwd = 1)

    for (i in seq_along(x)) {
        draw_ci(x[i], shift$lower_95_deg[i], shift$upper_95_deg[i])
    }

    points(x, shift$phase_shift_deg, pch = 16, cex = POINT_CEX)

    mtext("D", side = 3, adj = 0, line = 0.35, font = 2, cex = 2)
}

make_figure = function(configuration_L1, configuration_aligned, shift) {
    old_par = par(no.readonly = TRUE)
    on.exit(par(old_par), add = TRUE)

    par(mfrow = c(2, 2),
        mar = c(4.8, 5.2, 2.0, 1.0),
        oma = c(0, 0, 0, 0),
        mgp = c(3.0, 0.8, 0),
        tck = 0.015,
        las = 1,
        family = FONT_FAMILY,
        cex.axis = 1.2,
        cex.lab = 1.2)
    
    plot_panel_single_mode(configuration_L1, "walk", "A")
    plot_panel_single_mode(configuration_L1, "swim", "B")
    plot_panel_aligned(configuration_aligned)
    plot_panel_shift(shift)
}

configuration_L1 = read_data(CONFIG_L1_FILE)
configuration_aligned = read_data(CONFIG_ALIGNED_FILE)
shift = read_data(SHIFT_FILE)

validate_columns(configuration_L1, c("mode", "leg", "phase_deg_360"), CONFIG_L1_FILE)
validate_columns(configuration_aligned, c("mode", "leg", "phase_deg_360"), CONFIG_ALIGNED_FILE)
validate_columns(shift, c("leg", "phase_shift_deg", "lower_95_deg", "upper_95_deg"), SHIFT_FILE)

if (!all(c("walk", "swim") %in% unique(configuration_L1$mode))) {
    stop("The L1-reference configuration file must contain both walk and swim rows.")
}
if (!all(c("walk", "swim") %in% unique(configuration_aligned$mode))) {
    stop("The aligned configuration file must contain both walk and swim rows.")
}
if (!all(LEGS %in% configuration_L1$leg)) {
    stop("The L1-reference configuration file does not contain all six legs.")
}
if (!all(LEGS %in% configuration_aligned$leg)) {
    stop("The aligned configuration file does not contain all six legs.")
}
if (!all(LEGS %in% shift$leg)) {
    stop("The phase-shift file does not contain all six legs.")
}
configuration_L1$phase_deg_plot = phase_to_plot_range(configuration_L1$phase_deg_360)
configuration_aligned$phase_deg_plot = phase_to_plot_range(configuration_aligned$phase_deg_360)

cairo_pdf(filename = OUTPUT_PDF, width = 10.0, height = 9.6, family = FONT_FAMILY, onefile = TRUE)
make_figure(configuration_L1, configuration_aligned, shift)
dev.off()

png(filename = OUTPUT_PNG, width = 3000, height = 2880, res = 300, type = "cairo")
make_figure(configuration_L1, configuration_aligned, shift)
dev.off()

cat("\n============================================================\n")
cat("Six leg phase configuration figure\n")
cat("============================================================\n")
cat("Panel A: walking configuration with L1 fixed at 0 deg\n")
cat("Panel B: swimming configuration with L1 fixed at 0 deg\n")
cat("Panel C: aligned walking and swimming phase configurations\n")
cat("Panel D: walking-to-swimming phase change with 95% bootstrap CI\n\n")
cat("Input L1-reference configuration:\n", CONFIG_L1_FILE, "\n", sep = "")
cat("Input aligned configuration:\n", CONFIG_ALIGNED_FILE, "\n", sep = "")
cat("Input phase shift:\n", SHIFT_FILE, "\n", sep = "")
cat("Output PDF:\n", OUTPUT_PDF, "\n", sep = "")
cat("Output PNG:\n", OUTPUT_PNG, "\n", sep = "")
```
